# The Viral Protein Corona Directs Viral Pathogenesis and Amyloid Aggregation

**DOI:** 10.1101/246785

**Authors:** Kariem Ezzat, Maria Pernemalm, Sandra Pålsson, Thomas C. Roberts, Peter Järver, Aleksandra Dondalska, Burcu Bestas, Michal J. Sobkowiak, Bettina Levänen, Magnus Sköld, Elizabeth A. Thompson, Osama Saher, Otto K. Kari, Tatu Lajunen, Eva Sverremark Ekström, Caroline Nilsson, Yevheniia Ishchenko, Tarja Malm, Matthew J.A. Wood, Ultan F. Power, Sergej Masich, Anders Lindén, Johan K. Sandberg, Janne Lehtiö, Anna-Lena Spetz, Samir EL Andaloussi

## Abstract

Artificial nanoparticles accumulate a protein corona layer in biological fluids, which significantly influences their bioactivity. As nanosized obligate intracellular parasites, viruses share many biophysical properties with artificial nanoparticles in extracellular environments and here we show that respiratory syncytial virus (RSV) and herpes simplex virus 1 (HSV-1) accumulate a rich and distinctive protein corona in different biological fluids. Moreover, we show that corona pre-coating differentially affects viral infectivity and immune cell activation. Additionally, we demonstrate that viruses bind amyloidogenic peptides in their corona and catalyze amyloid formation via surface-assisted heterogeneous nucleation. Importantly, we show that HSV-1 catalyzes the aggregation of the amyloid beta peptide (Aβ_42_), a major constituent of amyloid plaques in Alzheimer’s disease, in-vitro and in animal models. Our results highlight the viral protein corona as an acquired structural layer that is critical for viral-host interactions and illustrate a mechanistic convergence between viral and amyloid pathologies.

## Introduction

The term “protein corona” refers to the layer of proteins that adhere to the surfaces of nanostructures when they encounter biological fluids. Nanoparticles adsorb biomolecules in biological fluids due to the high free energy of their surfaces^1^. The importance of the corona layer stems from the fact that it constitutes the actual surface of interaction with biological membranes or “what the cell sees” in the in-vivo context^2^. Hundreds of proteins have been identified to confer a distinct biological identity of nanoparticles in different microenvironments depending on their size, chemistry and surface modification (recently reviewed in^3^). These factors were found to be critical determinants of the biodistribution and pharmacodynamics of nanoparticles. On the other hand, the ability of the surfaces of nanoparticles to partially denature certain corona proteins exposing “cryptic epitopes” highlights the role of the protein corona in the toxicology of nanoparticles^4–6^. The formation of a protein corona is particularly important in the context of nanoparticle interaction with amyloidogenic peptides such as amyloid beta (Aβ_42_) and islet amyloid polypeptide (IAPP), which are associated Alzheimer’s disease (AD) and diabetes mellitus type 2 disease, respectively. Nanoparticles have been shown to catalyze amyloid formation via binding of amyloidogenic peptides in their corona, thereby increasing local peptide concentration and inducing conformational changes that facilitate fibril growth via a heterogenous nucleation mechanism^7,8^. This surface-assisted (heterogenous) nucleation has been demonstrated for several nanoparticles with different amyloidogenic peptides including IAPP and Aβ_42_ ^9,10^.

Here, we studied viruses in terms of their biophysical equivalence to synthetic nanoparticles in extracellular environments. As nanosized obligate intracellular parasites, viruses lack any metabolic activity outside the cell, and can thus be expected to interact with host factors in the microenvironment similar to artificial nanoparticles. In the current work, we used well-established techniques of nanotechnology to study the protein corona of respiratory syncytial virus (RSV) in comparison to herpes simplex virus type 1 (HSV-1) and synthetic liposomes. Additionally, we studied the interaction of both RSV and HSV-1 with amyloidogenic peptides.

RSV is an enveloped *Orthopneumovirus* with a diameter between 100 and 300 nm and a single stranded negative-sense RNA genome with 10 genes encoding 11 proteins^11^. It is a leading cause of acute lower respiratory tract infections in young children worldwide, causing up to an annual estimate of 34 million cases^12^. By the second year of life, nearly 90% of children get infected with RSV causing up to 196,000 yearly fatalities^13^. Reinfection with RSV occurs throughout life, usually with mild local symptoms in the upper airways^14^. However, reinfection in the elderly and immunocompromised individuals can lead to severe clinical disease in the lower airways. While natural infection leads to the production of neutralizing antibodies, the ability of these antibodies to protect from subsequent RSV infections appears to be incomplete ^15,16^. Neither a vaccine nor an antiviral therapy is yet available except for passive immunization using the anti-RSV monoclonal antibody palivizumab. Early vaccine trials using formalin-inactivated RSV led to enhanced disease with up to 80% of vaccinees being hospitalized and two dying following natural RSV infection ^14,16^. This led to the hypothesis that host immune responses play an important role in the pathophysiology of airway disease caused by RSV.

HSV-1 is an example of another virus with high prevalence, infecting nearly 70% of the human population ^17^. HSV-1 is a double-stranded DNA virus consisting of an icosahedral nucleocapsid surrounded by tegument and envelope with virion sizes ranging from 155 to 240 nm^18^. HSV-1 is a neurotropic virus that infects peripheral sensory neurons and establishes latency^19^. Latent HSV-1 is occasionally reactivated causing peripheral pathology and under certain circumstances it can migrate into the central nervous system causing herpes simplex encephalitis; the most common cause of sporadic fatal viral encephalitis^19^. In the context of the current work we focused on the presumptive role of HSV-1 in the pathology of AD. A number of risk factors have been associated with AD, including the E4 allele of the apolipoprotein E (Apo-E), diabetes, vascular pathology, neuroinflammation and infections^20^. Several recent studies have supported the theory of a significant role of HSV-1 in the disease ^21^. HSV- 1 DNA was found to be localized within amyloid plaques in AD patients and HSV-1 infection has been shown to promote neurotoxic Aβ accumulation in human neural cells and to the formation of Aβ deposits in the brains of infected mice^22,23^. Moreover, the presence of anti-HSV IgM antibodies, which indicate HSV reactivation, is correlated with a high risk of AD and antiherpetic treatment is correlated with a reduced risk of developing dementia^24,25^. Despite these correlations, the mechanism by which viruses induce amyloid aggregation, the major pathological hallmark of AD, is not known.

In the present study, we demonstrated that upon encountering different biological fluids, RSV accumulated extensive and distinct protein coronae compared to HSV-1 and synthetic liposomes. The various coronae were dependent on the biological fluid and exerted markedly different effects on RSV infectivity and capacity to activate monocyte-derived dendritic cells (moDCs). Moreover, upon interaction with an amyloidogenic peptide derived from IAPP, RSV accelerated the process of amyloid aggregation via surface-assisted heterogenous nucleation. This amyloid catalysis was also demonstrated for HSV-1 and the Aβ_42_ peptide in-vitro and in an AD animal model. Our findings highlight the importance of viral protein corona interactions for viral pathogenesis and provide a direct mechanistic link between viral and amyloid pathologies.

## Results

### Rich and unique protein coronae for RSV and HSV-1

Based on the extensive literature describing the significant role of corona factors in synthetic nanoparticle functionality, we used established techniques to answer questions regarding RSV pathogenicity^26^. Using proteomics, we assessed the RSV protein corona profiles in adult human plasma (HP), juvenile (6 months old infants tested RSV negative at the time of sample collection) human plasma (jHP), human bronchoalveolar lavage fluid (BALF) from healthy adults, rhesus macaque plasma (MP), and fetal bovine serum (FBS). These biological fluids represent different microenvironments encountered by the virus in terms of tissue tropism (HP vs. BALF), zoonosis (MP) and culturing conditions (FBS). The biological fluids were screened for antibodies against RSV using ELISA, and both adult HP and BALF contained high levels of anti-RSV IgG antibodies, unlike jHP, MP and FBS (Supplementary Fig. 1).

Viral stocks were produced in serum-free conditions to prevent initial contamination with FBS proteins. Virions produced under serum-free conditions were incubated with 10% v/v solutions of different biological fluids. Controls included non-infected cell medium representing the background cellular secretome, and synthetic lipid vesicles of a size comparable to RSV (200 nm) with positively or negatively charged surfaces. In addition, we compared the coronae of RSV to HSV-1 to probe for differences between different viruses of relatively similar size. After incubation for 1 h at 37 °C, the virions were re-harvested by centrifugation and washed twice before performing mass spectrometry-based proteomic analyses.

Assessment of the proteomic data by principle component analysis (PCA) showed that RSV and HSV-1 samples were well separated from one another and that the viral samples were well separated from the control samples with the replicates clustering together (Fig. 1 and Supplementary Fig. 2). Replicate clustering was confirmed via corresponding correlation plots for the samples in the PCA, which showed higher correlation coefficients between the replicates within the same corona condition than between different conditions (Supplementary Fig. 3). Notably, both RSV and HSV-1 possessed distinctive proteomic profiles depending on the biological fluid (Fig. 1c and Supplementary Fig. 2c). In addition, RSV protein corona profiles were different between HP and jHP (Supplementary Fig. 2d); however, the use of a different mass spectrometer for this experiment prevents direct comparison to the data shown in Fig. 1. The reproducibility of the corona preparations was further assessed by calculating coefficient of variation (CV) between the replicates for each sample type. The CVs ranged between 16-42% depending on sample type (average 28%, Supplementary Fig. 3d).

**Figure 1.**
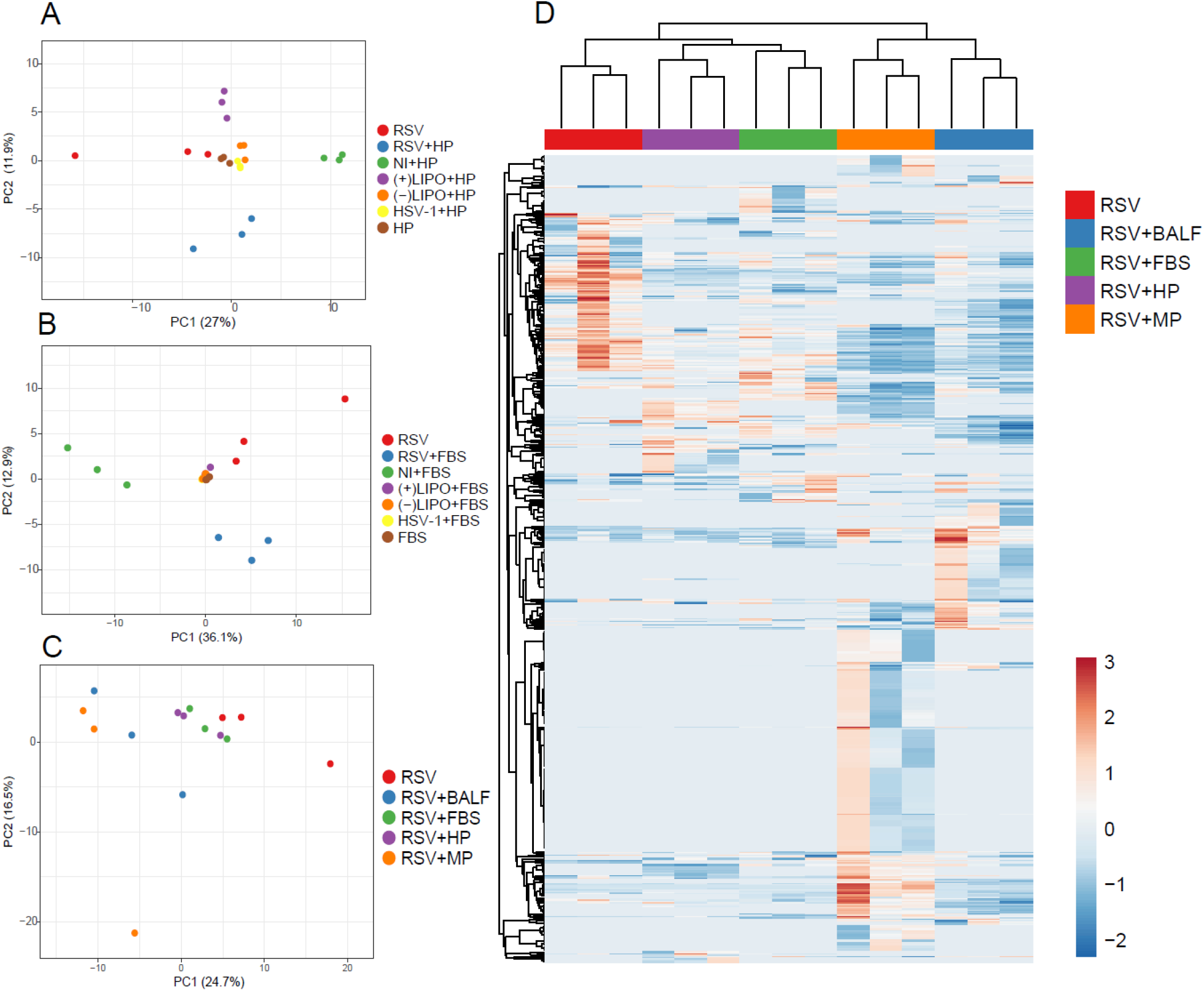
RSV accumulates a rich and distinctive protein corona in different biological fluids. (**a-c**) Principal component analyses (PCA) of the corona proteomic profiles of RSV, HSV-1 and controls. Triplicate samples were incubated with 10% solutions of each different biological fluid for 1h at 37 °C, then re-harvested, washed and finally analyzed by MS. Only proteins significantly detected (FDR 1%) in all three replicates in each condition were used. NI= Non-Infected supernatant, (-)Lipo = negatively charged lipid vesicles, 200 nm, (+)Lipo = positively charged lipid vesicles, 200 nm. (**a**) PCA comparing proteomic profiles in human plasma (HP) (**b**) PCA comparing proteomic profiles in fetal bovine serum (FBS) (**c**) PCA comparing the corona profiles of RSV in different biological fluids; HP, FBS, MP or BALF. (**d**) Heat map representing the viral corona fingerprints of RSV after incubation in different biological fluids. The three columns in the heatmap show three replicates. Only proteins significantly detected (FDR 1%) in all three replicates in each condition were used. Red and blue indicate higher and lower than the mean protein signal, respectively. Scale bars represent row Z-scores.

In order to determine if the most abundant corona factors were the most abundant proteins in the biological fluids, the crude fluids were also analyzed by mass spectrometry. The top 10 most abundant proteins identified are presented in Table 1. Similar to previous findings for nanoparticles^27^, our data showed that the most abundant proteins in the protein corona were not necessarily the most abundant in the biological fluids. This, together with the PCA analyses comparing RSV to HSV-1 and the controls indicated enrichment of particular corona factors depending on the surface properties of the virus. Moreover, proteomic data sets were visualized by heat map and hierarchical clustering revealing an extensive protein corona signature for viral samples in each biological fluid (Fig. 1d). While the serum-free viral particles contained some host factors that were incorporated during virus replication and budding from the cells, the virus was able to subsequently accumulate a different set of host factors which were dependent on the biological fluid. As such, a characteristic viral biological identity was associated with each biological fluid. The viral corona factors present on all three replicates in each biological fluid are listed in Supplementary Data 1. In addition, the raw proteomics data are available in Supplementary Data 2.

**Table 1.**
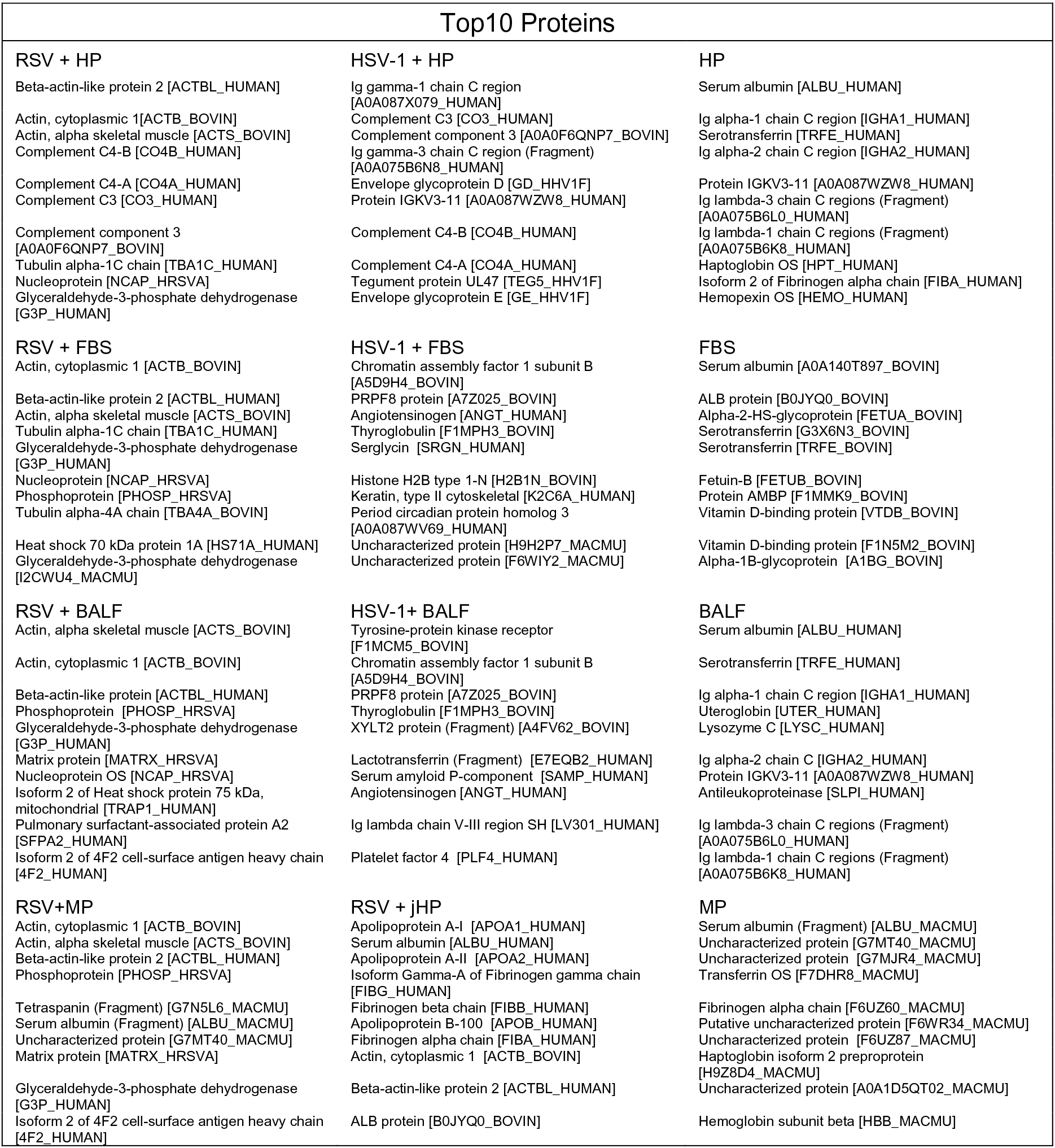
Top 10 proteins in the protein coronae of RSV, HSV-1 and crude biological fluids.

Additionally, we used transmission-electron microscopy (TEM) to visualize the viral protein corona. A layer of factors was observed interacting with the viral surface upon encounter with cell membranes, which was absent in serum free conditions (Fig. 2a). This demonstrated that RSV accumulated a layer of corona factors that are likely involved in cellular interactions. We also performed cryoimmuno electron microscopy using antibodies for certain proteins that were detected in the viral corona proteomic analysis. In serum-free conditions, we used an anti-RSV F-protein antibody. For adult HP and BALF coronae, we used anti-human IgG and anti-surfactant protein A (SP-A) antibodies, respectively. The bound antibodies were detected using secondary antibodies coated with 10 nm gold nanoparticles. As shown in Fig. 2b, corona factors were bound to the surface of RSV and labelled with the respective antibodies. To further confirm the binding of these factors, we performed western blot analysis on viral particles incubated with either HP or BALF. Several bands were detected for SP-A in the BALF corona samples and were completely lacking in the HP corona samples (Supplementary Fig. 4a). In addition to the main SP-A bands, several high molecular weight bands were also detected with the anti-SP-A antibody. This suggested that SP-A was forming complexes either on its own or with other corona factors. When stained with anti-human IgG antibodies, bands were detected in both coronae, but with higher intensity in the HP corona compared to the BALF corona (Supplementary Fig. 4b). The bands were normalized to the RSV G protein band which was used as a loading control (Supplementary Fig. 4c). Taken together, our data demonstrated that RSV acquired a differential protein corona layer depending on the biological fluid.

**Figure 2.**
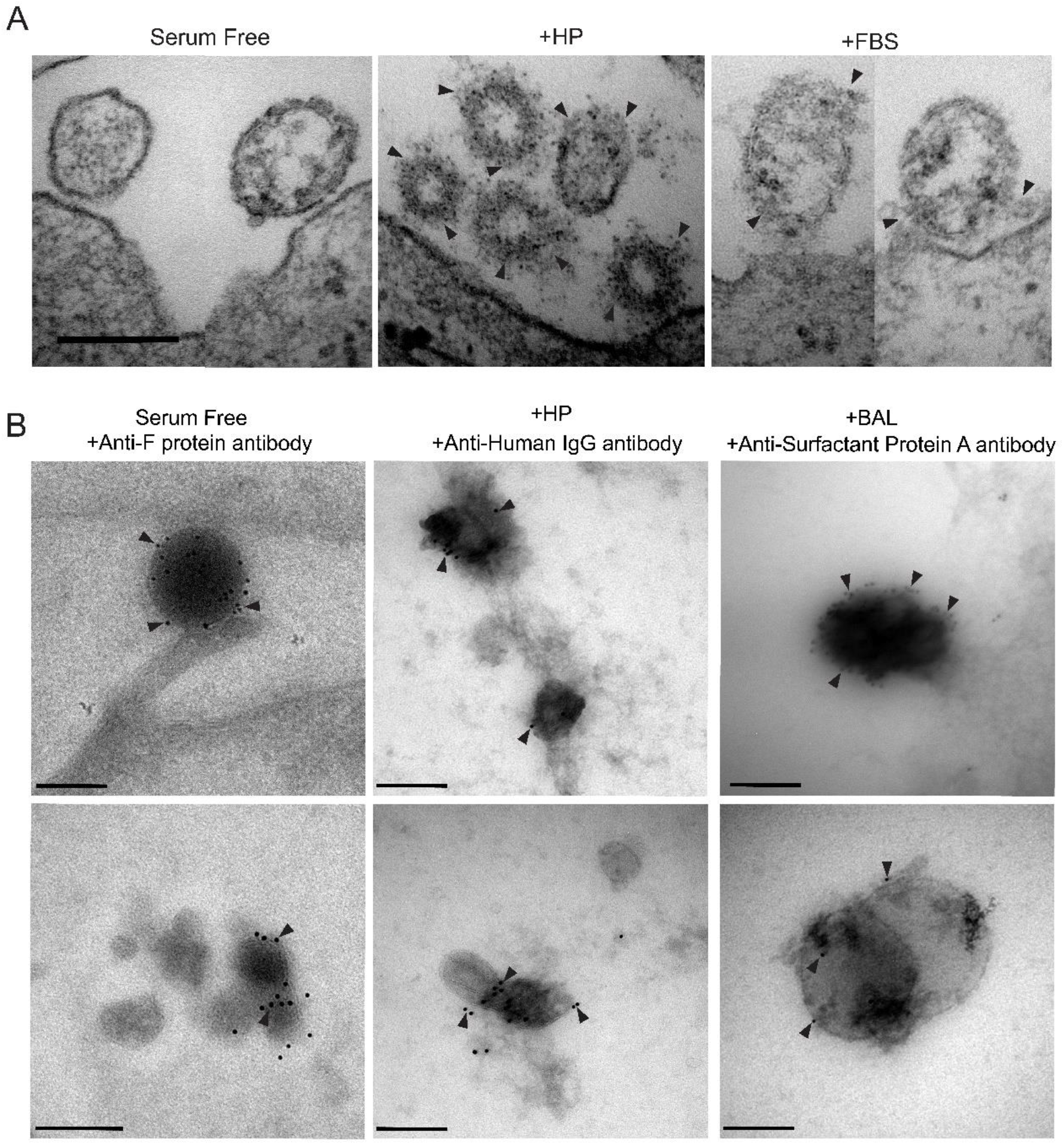
Corona factors bind to the viral surface. **(a)** Representative TEM images of HEp-2 cell sections (n=3) after incubation for 1h with RSV in either serum-free medium or medium with 50% v/v of different biological fluids. Compiled images of virions in close proximity to the cell-surface and black arrows point to bound protein corona **(b).** Representative cryoimmuno electron microscopy images of RSV incubated in serum-free conditions or with 50% v/v of different biological fluids (n=3) then labelled with anti-RSV F protein antibody, anti-human IgG, or anti-surfactant protein A antibody. Black arrows indicate gold labelling. Bar = 200 nm.

### Protein corona influences viral infectivity and moDC activation

To investigate if differential corona composition affects viral infectivity, virions produced under serum-free conditions were pre-incubated with different biological fluids before infection of HEp-2cells. The RSV-GFP virions were incubated with biological fluids at a protein concentration of 0.3 mg/ml (equivalent to 5% v/v) for 1 h at 37 °C then diluted 10 times in serum-free medium before infecting the cells at final MOI of 1. Corona pre-coating had a significant effect on viral infectivity as demonstrated by the differential frequency of GFP expressing cells quantified by flow cytometry (Fig. 3a-b and Supplementary Fig 5). HP corona pre-coating significantly reduced infectivity compared to serum-free conditions, while FBS and MP led to 5-6 fold enhancement in infectivity. The BALF corona also enhanced infectivity, but to a lower extent as compared to MP and FBS. Moreover, fluorescence microscopy revealed syncytia formation of cells infected with BALF, MP and FBS coronae (Supplementary Fig. 6). No significant differences in cell-toxicity were observed. jHP pre-coating, on the other hand, slightly enhanced infectivity, but it did not reach significance (Fig. 3b). This lack of inhibition compared to adult HP could be attributed to the lack of anti-RSV antibodies in the selected jHP (Supplementary Fig. 1). Furthermore, we investigated the effect of different coronae on the activation of human moDCs by quantifying the expression of the maturation marker CD86. Differentiated moDCs were infected by virions produced under serum-free conditions with different corona pre-coatings for 4h in serum-free conditions. The cells were then washed and incubated in serum-containing medium for 72 h prior to flow cytometry analyses. Only the BALF pre-coated virions were able to induce moDC activation and increase CD86 expression (Fig. 3c-f). Notably, adding BALF alone or corona-free RSV did not activate the cells, showing that it was the RSV-BALF corona complex that induced moDC activation. In addition, BALF pre-coating enhanced infectivity in moDCs as quantified by flow cytometry of GFP expression (Fig. 3e-f). Altogether, this shows that pre-incubation of virions produced under serum-free conditions in different biological fluids to allow a corona formation greatly affects infectivity and the ability to induce moDC activation in a biological fluid-dependent manner.

**Figure 3.**
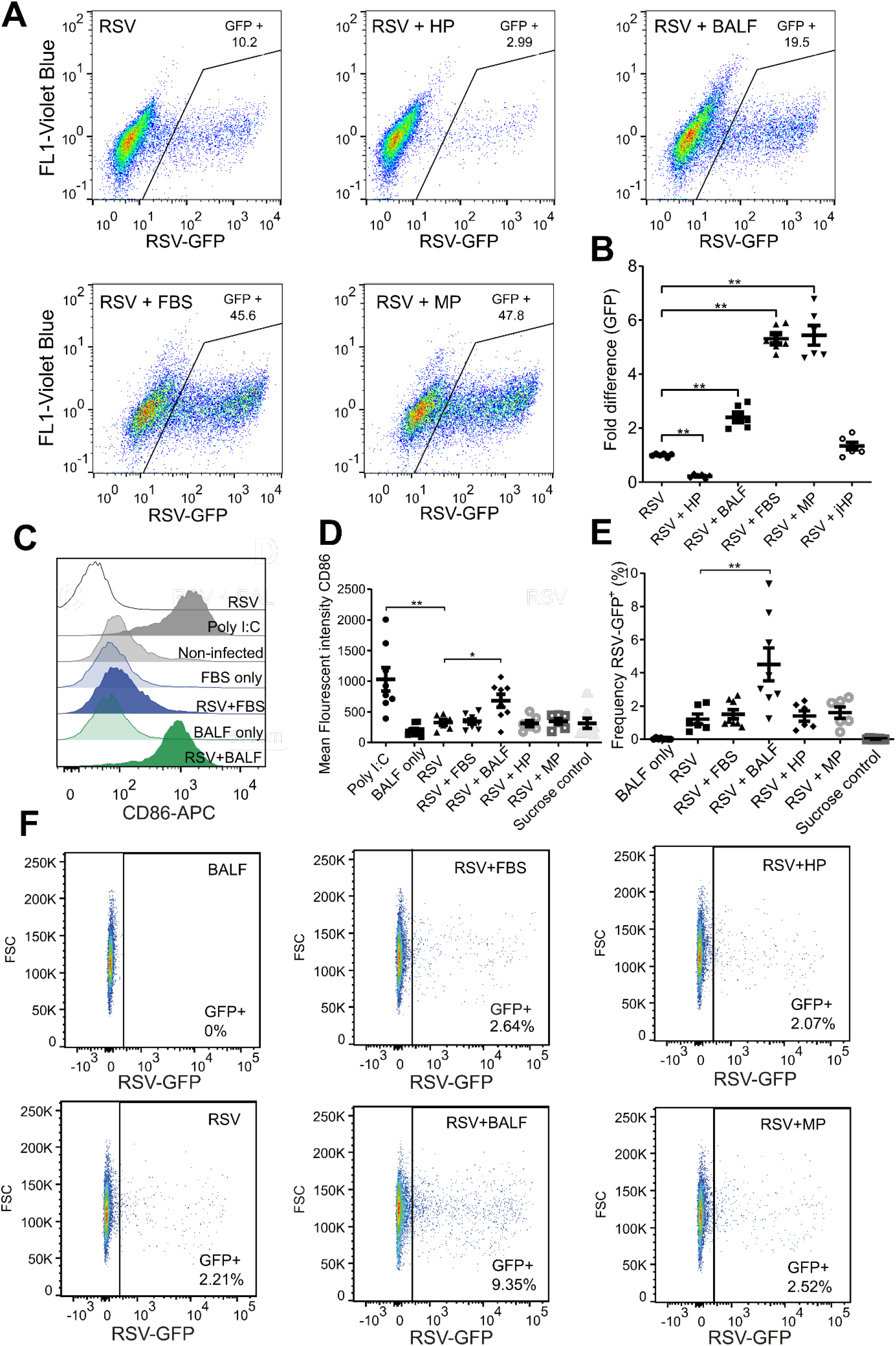
Different protein coronae affect RSV infectivity and moDC activation. **(a)** RSV virions produced under serum-free conditions were pre-coated with different coronae via pre-incubation with different biological fluids prior to addition to HEp-2 cells in serum-free medium (diluted 10x) at an MOI of 1. The frequencies of GFP expressing cells were quantified by flow cytometry 72 h post-infection. Gating strategy and uninfected control is shown in Supp Fig 5. Representative dot blot graphs are shown. **(b)** Flow cytometry quantification of the GFP positive-cells after RSV pre-coating with different coronae presented as fold-increase over RSV treatment in serum-free conditions (no corona). Means ±SEM of six replicates from two separate experiments are shown. Significant differences in comparison to RSV (no corona) were assessed by non-parametric Kruskal-Wallis unpaired test followed by Mann-Whitney test. P-value: ** *P* ≤ 0.01. (**c-f**) RSV virions produced under serum-free conditions were pre-coated with different coronae prior to the addition to primary moDCs in serum-free conditions at MOI of 20. CD86 expression and frequencies of RSV-GFP^+^ cells were quantified by flow cytometry 72 h post-infection (**C**) Representative histograms of CD86 expression in moDC are shown. (**d**) The mean fluorescent intensity (MFI) of CD86 expression in moDCs infected with RSV pre-coated with different coronae. Sucrose from the sucrose cushion used for RSV harvesting was also used as a control (**e**) The frequency of RSV-GFP expressing moDCs treated with RSV pre-coated with different coronae, and representative dot blot graphs are shown in (**f**). For moDCs, data are shown as MFI mean ± SEM for 6-8 individual donors from four separate experiments. Significant differences in comparison to RSV (no corona) were assessed by non-parametric Kruskal-Wallis unpaired test followed by Mann-Whitney test. P-value: ** *P* ≤ 0.01 and * *P* ≤ 0.05.

### Differential RSV corona composition

The set of factors that were only present in the HP corona and not in the other conditions comprised several antibodies and complement factors (Fig. 4a). Moreover, gene list enrichment analyses of HP corona revealed an enrichment of immunological gene ontology (GO) terms such as complement activation and humoral immune response (Fig. 4b). The HP corona proteomic profile was consistent with the observed inhibition of infectivity (neutralization), indicating that the corona characterization methodology is representative of the actual layer of host factors that surrounds the viral particle. Notably, despite having comparably high levels of virus-specific IgG antibodies, BALF imparted opposite effects on infectivity compared to HP (Fig. 3). At the proteomics level, GO analysis revealed that the viral BALF corona comprised a different set of factors that are less enriched in immunological components and more enriched in factors related to adhesion, anchoring, protein targeting to membrane, interspecies interaction, mutualism through parasitism, protein complex binding, and macromolecular complex binding (Fig.4). This was further supported by the western blot analysis where the corona of the BALF incubated virions was less enriched in human IgG compared to the viral HP corona (Supplementary Fig. 4).

**Figure 4.**
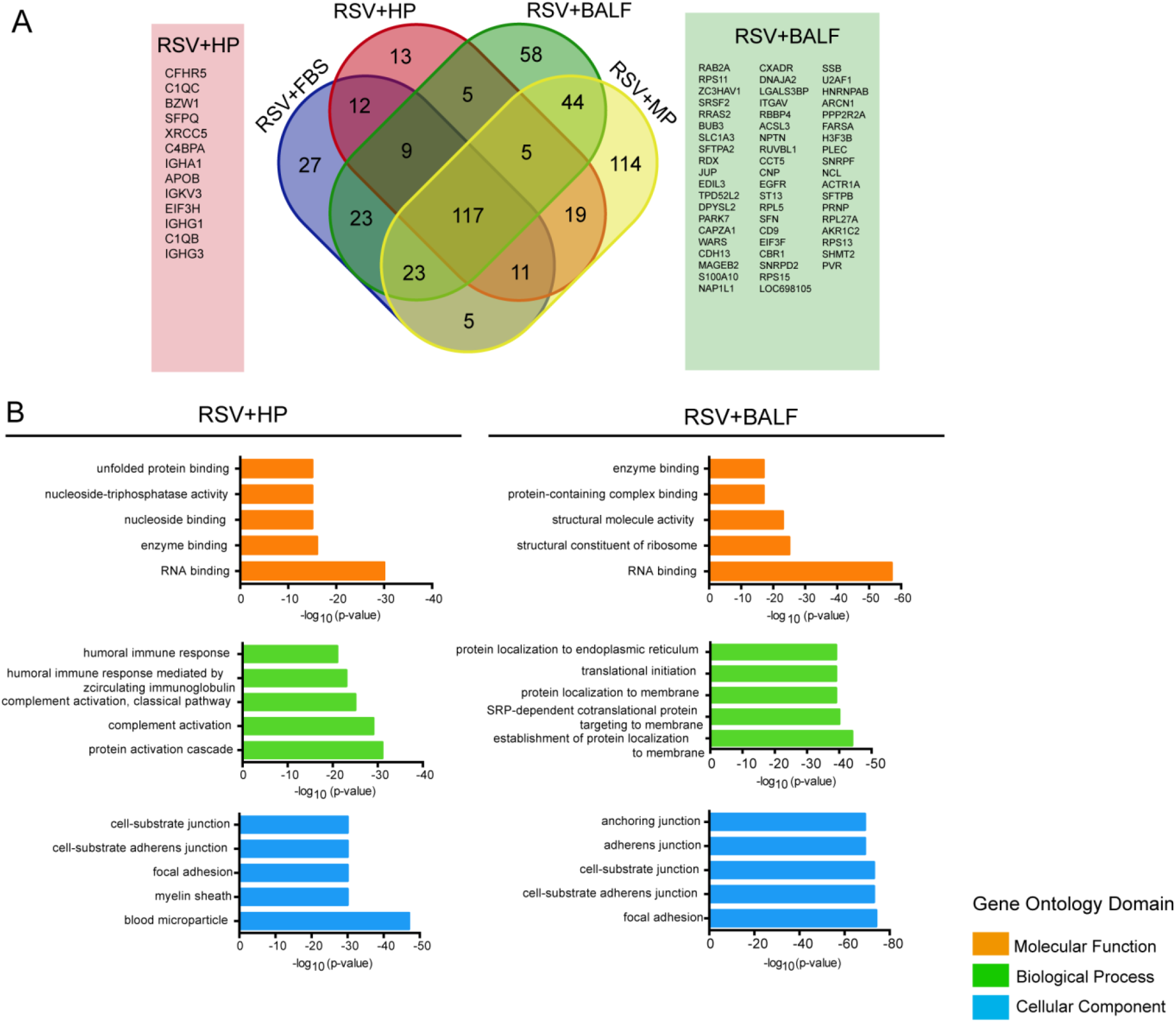
RSV corona proteomic representation and GO analysis in HP and BALF. (**a**) Venn-diagram showing the unique and overlapping protein populations from the RSV corona profiles in different biological fluids. Only proteins significantly detected (FDR 1%) in all three replicates in each condition were used. The unique factors in HP and BALF are shown. (**b**) Gene list enrichment analysis of the total RSV corona profile in BALF and HP groups. The top five enriched terms are shown in each GO domain.

On the other hand, anti-RSV IgG negative fluids (FBS and MP) enhanced viral infectivity in HEp-2 cells. The effect of both FBS and MP corona on enhancement of RSV infectivity was concentration-dependent as shown in Fig. 5a. While the FBS corona enhancement increased with concentration, the MP effect decreased after a protein concentration of 0.3 mg/ml was reached. This can be due to the increase of unbound corona factors that compete with bound factors for cellular receptors. Furthermore, we investigated the effect of corona pre-coating with FBS and MP on viral neutralization via palivizumab, which is a humanized monoclonal antibody directed against the F protein of RSV. We found that the enhancing effects of the coronae were completely lost in the presence of the antibody and that the antibody neutralization curves were very simillar in the presence and the absence of the viral corona (Fig. 5b). These data demostrated that high affinity antibodies were able to compete out corona factors to impart host protection. GO analysis revealed that FBS and MP coronae were also enriched in terms such as anchoring, adhesion, receptor binding, protein complex binding, unfolded protein binding, interspecies interaction, viral process, and mutualism through parasitism (Fig. 5c-d).

**Figure 5.**
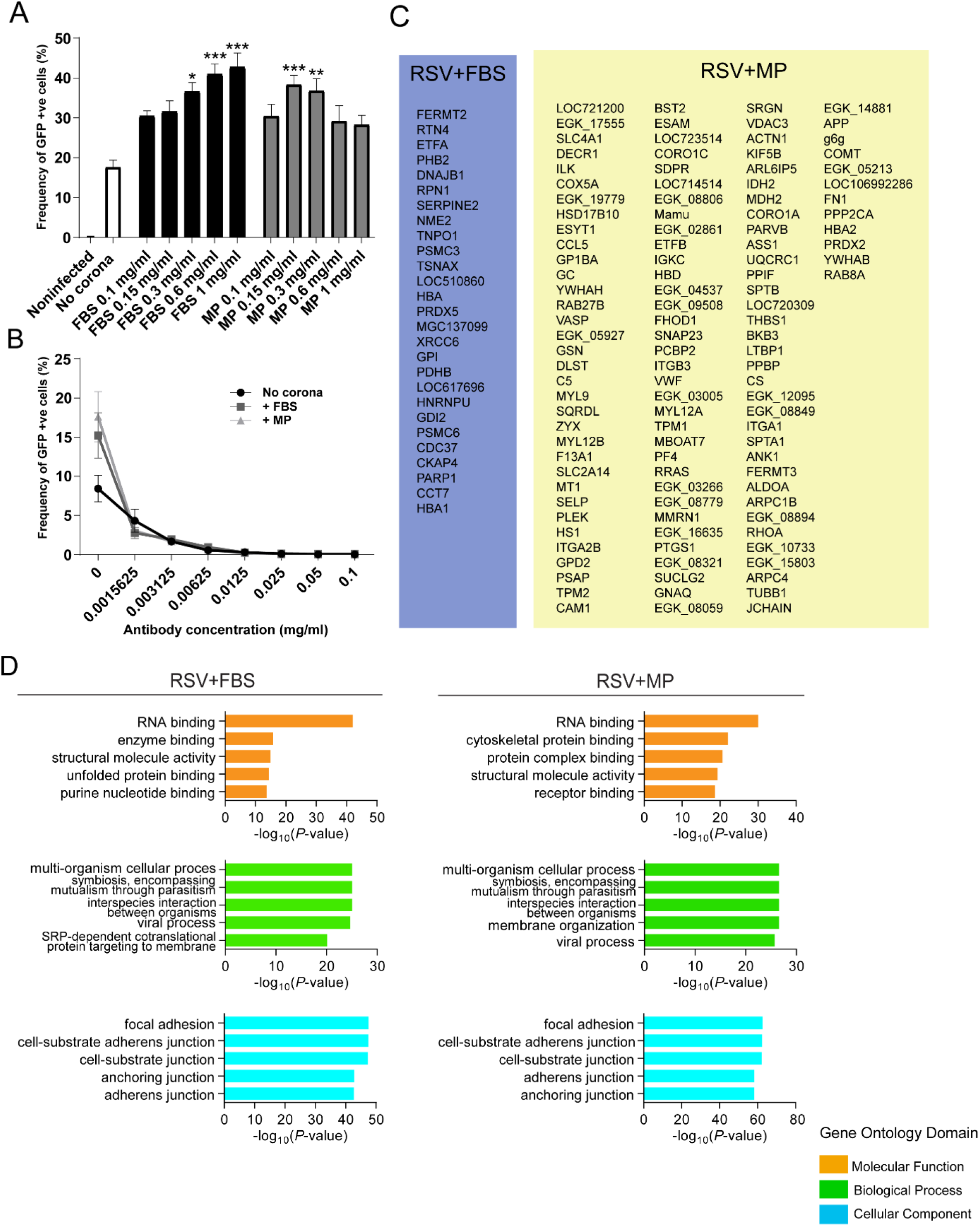
RSV protein corona in FBS and MP. (**a**) Serum-free produced RSV was pre-coated with different coronae via pre-incubation with different biological fluids prior to infection of HEp-2 cells in serum-free medium (diluted 10x) at an MOI of 1. The frequencies of GFP expressing cells were quantified by flow cytometry 72h post-infection. Means ±SEM of six replicates from two separate experiments are shown. Significant differences in comparison to RSV (no corona) were assessed by non-parametric Kruskal Wallis one-way ANOVA with Dunn’s multiple comparison test and are indicated by \**P* <.05, \*\**P* <.01 and \*\*\**P* <.001, respectively. (**b**) Neutralization curves with different concentrations of palivizumab (monoclonal antibody targeting RSV F-protein) of RSV with or without FBS and MP protein coronae at the protein concentration of 0.3 mg/ml protein. (**c**) The unique factors in FBS and MP from Fig. 4 are listed. (**d**) Gene list enrichment analysis of the total RSV corona profile in FBS and MP groups. Only proteins significantly detected (FDR 1%) in all three replicates in each condition were used. The top five enriched terms are shown in each GO domain.

### RSV catalyzes amyloid aggregation

Since nanoparticles are known to bind amyloidogenic peptides in their coronae leading to induction of amyloid aggregation via a heterogenous nucleation mechanism, we next investigated whether viruses are also capable of this particular corona interaction. We investigated the interaction of RSV with a model amyloidogenic peptide (NNFGAIL) derived from the IAPP. We traced the ability of RSV to accelerate amyloid kinetics using the well-established thioflavin T (ThT) based methodology. The ThT dye changes its fluorescence emission spectrum upon binding to amyloid fibrils, and plotting relative changes in the fluorescence intensity against time illustrates the kinetics of the amyloid formation process^28^. Using the ThT assay, we found that the presence of RSV particles significantly accelerated amyloid formation of NNFGAIL compared to non-infected cell supernatant demonstrating that RSV acted as a catalytic surface for amyloid aggregation (Fig. 6a). As a control, we compared the kinetic curves of the virus-containing medium vs. virus-free medium upon incubation with ThT alone without a peptide. As shown in Supplementary Fig 7a, the curves were very similar, indicating that the relative changes that we observed in the presence of amyloidogenic peptides are not due to unspecific interaction of the virions with the ThT dye. Additionally, extensive fiber networks were observed with TEM within 100 min. of incubation with RSV (Fig. 6b) and some virions were located at the tip of these fibers (Fig. 6c). Moreover, RSV catalytic activity was more efficient than lipid vesicles of similar size; however, it was dramatically reduced in the presence of 5% FBS, indicating a competition between the peptide and other corona factors for the viral surface (Fig. 6d). On the other hand, RSV failed to catalyze the amyloid aggregation of GNNQQNY peptide, which is derived from yeast prion protein (Fig. 6e). This, in turn, indicated that such interactions are not universal to all amyloidogenic peptides but display some selectivity depending on the virus/peptide pair.

**Figure 6.**
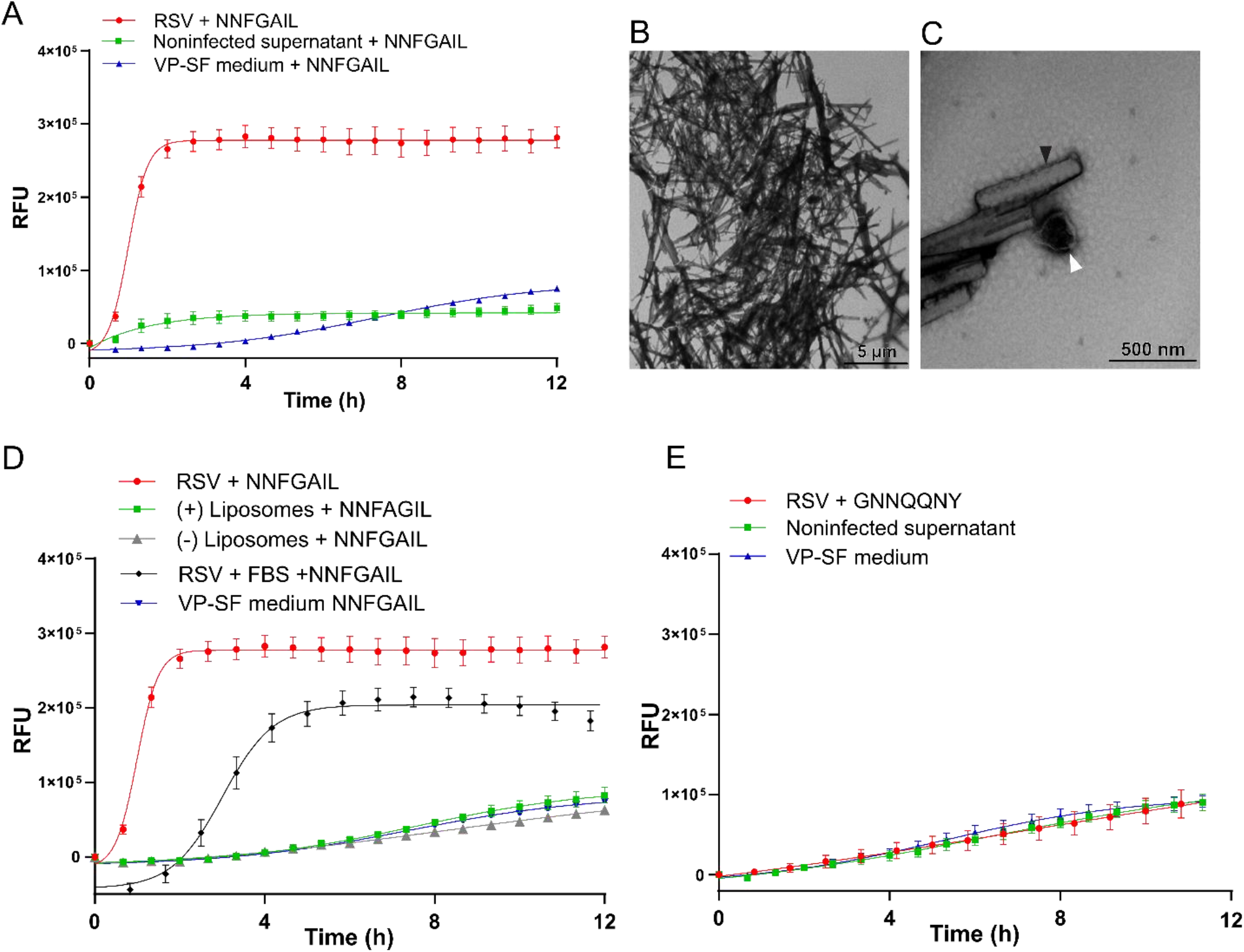
RSV accelerates the kinetics of amyloid formation. (**a**) NNFGAIL peptide incubated with ThT solution and RSV (3 × 10^8^ genome copy/ml), non-infected supernatant or serum-free viral production medium (VP-SF medium). ThT fluorescence was measured at 440 nm excitation and 480 nm emission over 12 h at 37°C. Means ±SEM of six replicates from two separate experiments are shown. (**b-c**) Negatively stained TEM images of RSV incubated with 1 mM NFGAIL for 100 mins. at 37°C. (**b**) Fibrillar tangles are shown (**c**) RSV virion shown at the base of an amyloid fiber. White arrows indicate viral particle and black arrows indicate fibrillar structures. (**d**) NNFGAIL peptide incubated and monitored at similar conditions in the presence or absence of 5% FBS, lipid vesicles (positively and negatively charged, dimeter= 200 nm, concentration = 1 × 10^10^ particles/ml) or VP-SF medium. Means ±SEM of six replicates from two separate experiments are shown. (**e**) GNNQQNY peptide incubated with RSV, non-infected supernatant or VP-SF medium. Means ±SEM of six replicates from two separate experiments are shown.

### HSV-1 catalyzes Aβ_42_ amyloid aggregation in-vitro and in-vivo

The finding that viral surfaces could serve as catalysts for amyloid formation warranted further investigation and confirmation using another virus/peptide system. To this end, we investigated HSV-1 and the Aβ_42_ peptide, whose aggregation is a major hallmark of AD. Recently, there has been an increasing body of reports suggestive of a correlation between HSV-1 infection and AD, reviewed in ^21^. However, evidence of a direct role of HSV-1 in the process of amyloid nucleation and subsequent fibril growth is currently lacking.

We found that HSV-1 significantly accelerated amyloid formation of Aβ_42_ compared to non-infected cell supernatant (Fig. 7a). As a control, and similar to the RSV/NNFGAIL system, both the virus-containing medium and virus-free medium produced similar curves upon incubation with ThT alone without peptide (Supplementary Fig. 7b). The catalytic activity was reduced by the presence of FBS, also indicating a competition at the viral surface (Fig 7b). In addition, HSV-1 was more efficient than liposomes in accelerating the amyloid aggregation kinetics (Fig. 7c). Furthermore, the propensity of HSV-1 mediated amyloid catalysis was higher for the more amyloidogenic Aβ_42_ peptide compared to the shorter, less amyloidogenic Aβ_40_ peptide (Fig. 7d). Amyloid induction was further confirmed by TEM demonstrating fibril formation within 100 min. of incubation with viral particles (Fig. 7e-h). Amyloid protofilaments and fibrils at different stages of elongation were observed interacting with the viral surface. Fig. 7e and f show multiple fibrillary structures emerging from one viral particle suggesting that a nucleation mechanism was taking place on the surface, thereby sparking fibril elongation. Viral particles also interacted with fibrillar structures that are part of an extensive network of fibers as shown in Fig. 7g and h. Importantly, to demonstrate the in-vivo relevance of our mechanistic findings, we intracranially infected transgenic 5XFAD mice with HSV-1. The 5XFAD mouse is a widely-used AD model as it recapitulates many AD phenotypes with rapid onset of Aβ_42_ aggregation that spreads to the hippocampus and cortex by six months of age^29^. We observed a significant increase in Aβ_42_ accumulation in the hippocampi and cortices of HSV-1 infected mice 48h post infection (Fig. 7i) in comparison with animals injected with non-infected supernatant. Representative images of the amyloid staining demonstrate the dramatic difference in amyloid accumulation between infected and non-infected animals (Fig. 7j). These results validated our biophysical findings using the ThT assay and demonstrated that the viral corona-driven catalysis of amyloid aggregation could take place in in-vivo situations.

**Figure 7.**
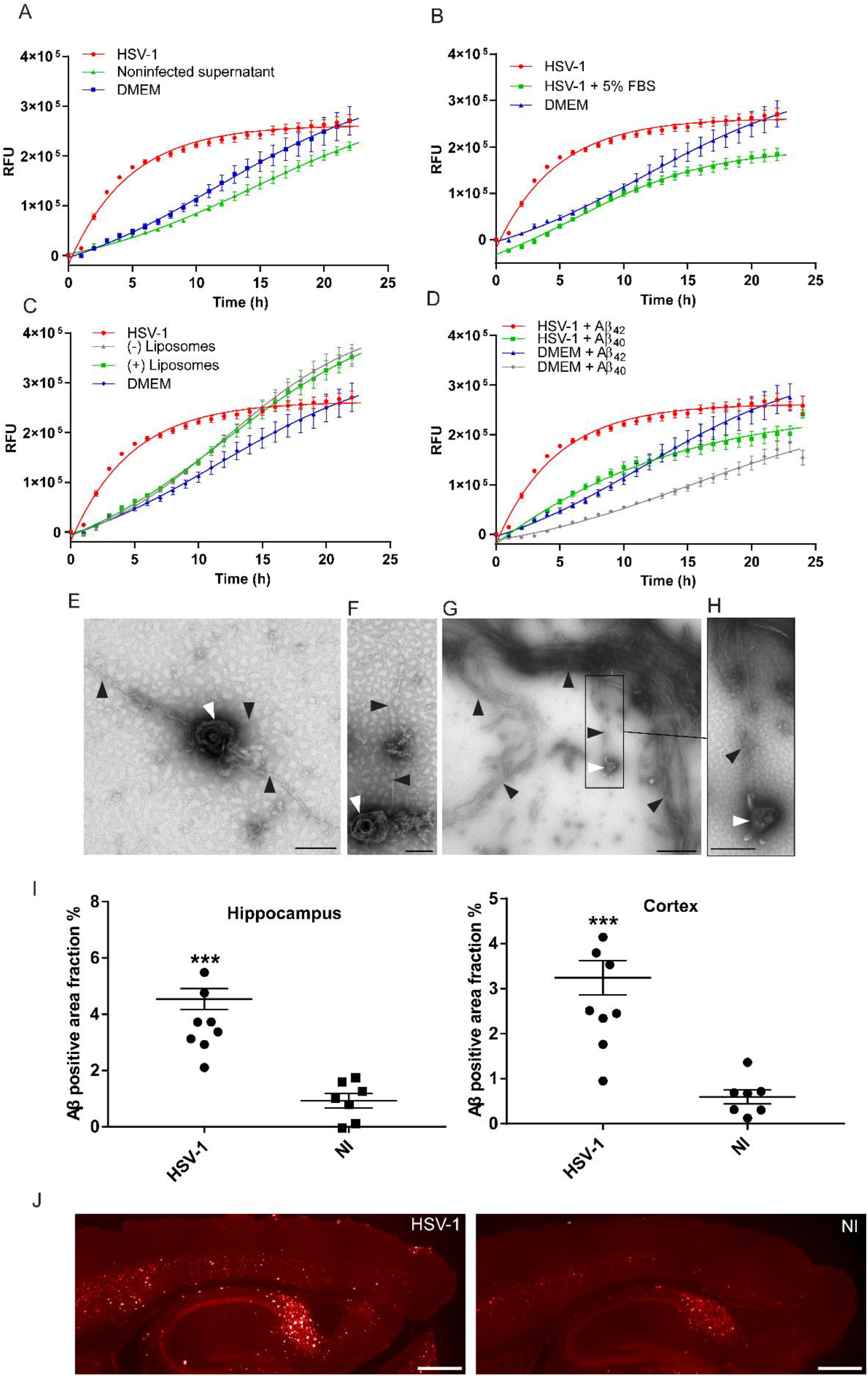
HSV-1 accelerates the kinetics of Aβ_42_ fibrillation in-vitro and in-vivo. (**a**) Aβ_42_ peptide was incubated with ThT solution and HSV-1 (2 × 10^8^ PFU/ml), non-infected supernatant or DMEM serum free growth medium. ThT fluorescence was measured at 440 nm excitation and 480 nm emission over 24 h at 37 °C. (**b**) Aβ_42_ incubated with HSV-1 in the presence or absence of 5% FBS. (**c**) Aβ_42_ incubated with either HSV-1 or lipid vesicles (positively or negatively charged, dimeter= 200 nm, concentration= 1 × 10^10^ particles/ml) incubated with Aβ_42_ peptide. (**d**) HSV-1 and DMEM serum free growth medium incubated with Aβ_42_ or Aβ_40_ peptides. For all the curves, means ±SEM of six replicates from two separate experiments are shown. (**e-h**) Negatively stained TEM images of HSV-1 incubated with 50 µM Aβ_42_ for 100 min. at 37 °C. White arrows indicate viral particles and black arrows indicate fibrillar structures. (**e**) Three protofilaments/fibrils stemming from one viral particle, bar = 200 nm. (**f**) Two protofilaments/fibrils stemming from a viral particle interacting with and aggregated structure, bar = 100 nm. (**g**) A viral particle interacting with protofilaments/fibrils which are connected to extensive fibrillar structures, bar =1 µm. (**h**) A rectangular close-up of viral interaction, bar = 500 nm. (**i-j**) Three-month-old transgenic 5XFAD mice were intracranially injected with HSV-1 or non-infected supernatant (NI). The brains were cryosectioned and stained using primary anti-body specific to isoforms Aβ 37-42 and then visualized by fluorescent Alexa 568 secondary antibody. (**i**) hippocampal and cortical Aβ immunoreactivities were quantified from 6 stained sections and are presented as positive area fraction percent. N=8 in the HSV-1 group and N=7 in the NI group. Significant differences were assessed by non-parametric Kruskal-Wallis unpaired test followed by Mann-Whitney test and are indicated by \*\*\**P* <.001. Data are shown as mean positive area fraction percent ± SEM from two separate experiments. (**j**) Representative images of the Aβ staining from HSV-1 vs. NI mice. Bar=500µm.

## Discussion

Viruses rely on the cellular machinery of the host for replication, production of viral proteins, assembly and export out of the cell. However, outside cells, viruses share many biophysical properties with artificial nanoparticles. Based on this biophysical equivalence, we hypothesized that viruses accumulate a rich and selective protein corona layer in the extracellular environment similar to nanoparticles. Examples of particular host factors that bind to viral surfaces have previously been described. For example, lipoproteins such as apolipoprotein E (Apo-E) were shown to be essential for hepatitis C virus (HCV) infection ^30^. Furthermore, coagulation factors such as factor X were shown to directly bind adenovirus leading to liver tropism ^31,32^. Other soluble components such as Gas6 were shown to contribute to the infectivity of lentiviral vectors even when pseudotyped with multiple types of envelope proteins ^33^. Furthermore, soluble heparin sulfonated proteoglycans (HSPGs) were shown to enhance the infectivity of human papillomavirus (HPV) ^34^. Interestingly, amyloids derived from prostatic acidic phosphatase (PAP) in semen were shown to enhance HIV-1 infectivity by several orders of magnitude ^35^. Additionally, plant viral nanoparticles that are used for drug delivery were shown to possess a protein corona that affects their uptake and biodistribution ^36^. However, to our knowledge, there is no previous work that has characterized the protein corona of an infectious animal virus in different biological fluids and studied its effect on viral pathogenicity.

In this work, we used proteomics to study viral protein corona enrichment in different biological fluids. We investigated the RSV protein coronae in biological fluids that are relevant to viral tropism, zoonosis and culturing conditions. As RSV is a virus tropic to the respiratory tract, we compared the viral corona in human BALF versus HP from adult healthy volunteers and jHP from RSV-seronegative 6-month-old infants. We also investigated the RSV corona in plasma from rhesus macaques, which are used as models in RSV studies and vaccine development^37^. Moreover, we studied the corona in FBS, which is the most commonly used cell growth supplement for viral production and in-vitro studies. We compared the RSV corona to HSV-1 and lipid vesicles of similar size with positively or negatively charged surfaces as well as the biological fluids per se. PCA, quantitative intra-sample variability analysis and correlation matrices demonstrated that the RSV corona profiles were well separated from that of HSV-1 and lipid vesicles (Fig. 1, Supplementary Fig. 2 and 3). Intra-sample variability could arise in the process of sample preparation during incubation, washing and digestion or during mass spectroscopy-based detection^38^. Notably, while the viral corona profile was dependent on the biological fluid, it was not a mere reflection of the most abundant proteins, as demonstrated earlier for nanoparticles^27^ and as shown in the Table 1 of the top 10 proteins present in the viral coronae in comparison to crude biological fluids. The viral protein corona was visualized using TEM, which revealed a layer surrounding the viral surface involved in cellular interactions (Fig. 2a). Using cryoimmuno electron microscopy and western blot, we could detect specific proteins identified from the proteomic analysis (human IgG in HP and SP-A in BALF) associated with RSV upon incubation with HP or BALF, respectively (Fig. 2b and Supplementary Fig. 4). Functionally, the coronae from the different biological fluids enhanced RSV infectivity except for the HP corona, which completely neutralized the virus (Fig. 3). Taken together, using several complementary methods, our data demonstrated the specific enrichment of differential corona profiles on viral surfaces. As such, the protein corona represents an initial phase of viral-host interactions that precedes cellular interaction and affects subsequent infectivity. Unlike the viral genome-coded surface proteins, the viral protein corona is an acquired structural layer that is dependent on the viral microenvironment resulting in different viral identities based on the target tissue and the target organism. Moreover, as the corona layer is a rich and complex layer, the final biological effect is expected to be dependent on a multitude of corona factors rather than a single protein.

Proteomic analysis of the HP corona revealed that it was enriched in antibodies and complement factors, which could explain the neutralization effect (Fig. 3 and 4). Other factors of documented immunological functions such as fibrinogen^39^, properdin^40^, annexin A1^41^, protein S100^42^ and vimentin^43^ were also detected in the HP corona (Supplementary Data 1). These factors could be parts of an immunological corona that enables efficient viral neutralization and/or immune system modulation. On the other hand, jHP, which lacked the anti-RSV antibodies, failed to neutralize the virus and the jHP corona was less enriched in immunological factors compared to the HP corona (Table. 1 and Fig. 3). Interestingly, the BALF corona enhanced viral infectivity in both HEp-2 cells and moDCs despite having equally high anti-RSV IgG antibody titers as compared to HP according to ELISA analysis (Supplementary Fig. 1). The proteomic and GO analysis showed that the BALF corona was less enriched in immunological factors (Fig. 4). Additionally, less IgG was detected in the BALF corona in the western blot analysis (Supplementary Fig. 4). The failure of BALF to neutralize the virus despite being rich in anti-RSV antibodies suggested that the differential protein corona profile could affect antibody binding in BALF conditions. This might explain the recurrence of pulmonary RSV infection even in individuals with high IgG titers^15^. An alternative explanation is that the BALF anti-RSV antibodies are of lower affinity than their HP counterparts and are thus less able to compete out other corona factors that enhance viral infectivity. It remains to be elucidated, which avidity is required in order to compete out the corona layer. The list of proteins that were uniquely detected in the BALF corona included pulmonary surfactants which are known to enhance RSV infectivity ^44^, nucleolin and EGFR, which were shown to be important for RSV cell entry^45,46^, adhesion molecules (such as tetraspanin, neuroplastin, integrin and cadherin), coxsackievirus/adenovirus receptor, poliovirus receptor and zinc finger CCCH-type antiviral protein 1 (Fig. 4). Notably, pre-incubation of RSV with BALF was the only condition that induced moDC activation (Fig. 3). Importantly, it was not the combination of the proteins in the biological fluids or the virus per se that affected the outcome, but rather the viral-corona complex, as shown by the lack of moDC activation by BALF or RSV alone. Moreover, and in accordance with the documented role of corona factors in nanoparticle immunogenicity^47^, our results demonstrated the role of the acquired corona layer in virus-induced immune activation that might contribute to disease pathophysiology. It can be speculated that the viral corona factors are part of the pathogen associated molecular patterns (PAMPS), which are recognized by the innate immune system. It also suggests that corona factors need to be considered in the design of vaccines and adjuvants for efficient stimulation of the immune system.

Additional biological fluids investigated in our study included FBS and MP and they both enhanced the infectivity of RSV in a concentration dependent manner (Fig. 5a). Artificially adding anti-RSV antibodies (palivizumab) successfully neutralized the virus even in these corona conditions, indicating successful out-competition by the antibody (Fig. 5b). GO analysis suggested a role of FBS and MP corona factors in interspecies interaction, mutualism, viral process, protein complex binding, unfolded protein binding, anchoring and adhesion (Fig. 5 c-d). Factors uniquely present in the FBS corona that may contribute to these effects include C4b-binding protein alpha-like which is a complement inhibitor, isoform 2 of fermitin family homolog 2 which binds to membranes enriched in phosphoinositides and enhances integrin-mediated adhesion ^48^ and hepatitis A virus cellular receptor 1 (Fig. 5c). In the MP corona, several adhesion proteins were found including: fibronectin isoform 4 preproprotein, endothelial cell-selective adhesion molecule, fermitin family homolog 3 short form and zyxin. The MP corona also contained receptor ligands such as: transferrin and C-X-C motif chemokine together with tetherin (isoform 2 of Bone marrow stromal antigen 2), which possesses antiviral properties^49^. Taken together, this illustrated that the observed functional effects of the viral protein corona were most likely mediated by a combination of factors that are enriched on the viral surface and not by a single factor. Since many viruses bind to several receptors and co-receptors, such interactions might be taking place in a multivalent manner ^50^. In addition, our results also highlight that the viral protein corona has to be taken into consideration in relation to zoonosis and applications involving viral propagation in-vitro.

We then investigated whether viral corona interactions with host factors involve surface-assisted nucleation of amyloids. Nanoparticles have been shown to act as catalytic surfaces that facilitate heterogenous nucleation of amyloid fibrils via binding, concentrating and enabling conformational changes of amyloidogenic peptides^7,51,52^. Similar to what have been reported with nanoparticles, we found that RSV accelerated the kinetics of amyloid aggregation of a model amyloidogenic peptide derived from IAPP (NNFGAIL). This demonstrated that viral particles are also capable of amyloid catalysis via surface-assisted heterogenous nucleation (Fig. 6). In order to investigate if this catalytic mechanism extends to other virus-amyloid pairs, we evaluated whether HSV-1 could accelerate the amyloid kinetics of Aβ_42,_ which is implicated in AD. Several recent studies have suggested HSV-1 involvement in AD^53^. Herein, we found that HSV-1 accelerated the kinetics of amyloid aggregation of Aβ_42_ and to a lesser extent the aggregation of Aβ_40_ (Fig. 7). HSV-1 was more efficient than lipid vesicles in amyloid catalysis and this efficiency decreased in the presence of FBS demonstrating a competition with other corona factors on the viral surface. Additionally, TEM demonstrated an interaction between amyloid fibrils at different stages of maturation and the viral surface via early protofibrillar intermediates, which we speculate represent the surface-assisted nucleation process. Importantly, infecting 5XFAD AD animal models with HSV-1 lead to increased accumulation of Aβ_42_ plaques as evident by immunohistochemical analysis (Fig. 7), demonstrating that the viral corona-driven catalysis can take place in-vivo.

Taken together, our data on RSV and HSV-1 demonstrated that viruses can physically act as nano-surfaces capable of catalyzing amyloid nucleation and leading to accelerated fibril formation. While our results do not prove disease causality, they present mechanistic explanation of the clinical and experimental correlations drawn between HSV- 1 and AD, which requires further investigation. Interestingly, Apo-E, which is a well-known risk factor for AD, was enriched in the HSV-1 corona, suggesting even further disease links (Supplementary Data 1).

Several recent studies have suggested that Aβ_42_ is an antimicrobial peptide that aggregates to sequester pathogens^54–57^. Here, we present an alternative but not mutually exclusive explanation. Our corona-driven hypothesis suggests that the virus interacts with extracellular amyloidogenic peptides as part of the pathogenesis, and the bound peptides do not necessarily possess an immunological and/or an antimicrobial function. This is further corroborated by studies showing that amyloids can enhance the infectivity for viruses such as HIV and HSV^35,58^, and the fact that amyloid precursor protein (APP) knock-out animals are not more susceptible for infections compared to wild-type animals^59^. From this perspective, virus surface-assisted nucleation might have evolved as a mechanism by which viruses modulate the host’s extracellular environment by catalyzing the phase transition of certain peptides from soluble to insoluble forms; a phase change that could lead to toxic gain or loss of function or both. In addition, the viral surface-assisted nucleation mechanism could be extended to other viruses correlated with neurodegenerative pathology such as HIV and HIV-associated neurocognitive disorder (HAND)^60^ and influenza virus and post-encephalitic parkinsonism ^61^ and others. Furthermore, the phenomenon of amyloid polymorphism^62^ might in part be related to the different nucleation mechanisms, where we hypothesize that the viral-catalyzed heterogeneously nucleated amyloids would be polymorphically distinct from homogeneously-nucleated amyloids that result from aberrant protein expression/overexpression. This hypothesis can be tested in future studies on the Aβ polymorphism in familial and sporadic Alzheimer’s disease ^63^, and whether this structural polymorphism can be traced back to different nucleation mechanisms related to different etiologies. Finally, the implication that viruses are capable of inducing conformational changes in bound host factors leading to exposure of cryptic epitopes may prove important for better understanding the correlation between viruses and autoimmune diseases.

To conclude, the current work was based on the biophysical equivalence between viruses and artificial nanoparticles in extracellular environments. We here demonstrated that nanotechnological concepts such as the protein corona and surface-assisted nucleation can be extended to infectious viruses. We showed that viral protein corona accumulation and amyloid catalysis are two aspects of the same phenomenon, namely viral surface interaction with extracellular host proteins. This phenomenon leads to the modulation of how viruses interact with cells and/or the induction of conformational changes in bound proteins that leads to accelerated amyloid aggregation. These findings highlight the potentially critical role of viral extracellular interactions in viral infectivity and in relation to extracellular protein pathology.

## Methods

### Viral and cell culture

HEp-2 cells, a human laryngeal carcinoma cell line, were used for RSV culture and infectivity experiments. Preparation of RSV A2-GFP stocks was performed using VP-SFM serum-free, ultra-low protein medium containing no proteins, peptides, or other components of animal or human origin (ThermoFisher, USA). HEp-2 cells were initially seeded in growth medium (DMEM medium with 5% FBS (ThermoFisher, USA), 1% Penicillin/Streptomycin (ThermoFisher, USA) and 0.01M HEPES (SigmaAldrich, Germany) until they reached approximately 70-80% confluency. At the day of infection, the cells were washed twice with warm PBS and the medium was replaced with VP-SFM containing 50 µg/ml gentamycin. Cells were infected at MOI of 4 and incubated for 5-6 days until >90% GFP expression was visible with fluorescence microscopy. Cells were then scraped, vortexed thoroughly, sonicated for 10 min., vortexed thoroughly one more time and spun at 1000 g for 5 min. The supernatant was transferred to new tubes and used freshly for the proteomics and amyloid interaction experiments. For long-term storage, infectivity and moDC activation experiments the supernatant was adjusted with MgSO_4_ and HEPES (SigmaAldrich, Germany) solutions to a final concentration of 0.1M MgSO_4_ and 50 mM HEPES. RSV was then concentrated using a 1.45 M sucrose cushions in the following buffer (1 M MgSO4; 50 mM Hepes, pH 7.5; 150 mM NaC1) according to ^64^ via centrifugation for 4 h at 7500 g at 4 °C. The viral concentrated layer was then harvested from interface between the cushion and the supernatant, aliquoted and immediately frozen at −80 °C. Quantifications of viruses were measured using real-time qPCR. Purification of viral RNA was performed using the QIAamp® Viral RNA extraction kit (QIAGEN) and extracted RNA was stored at −80 °C. Viral titers were determined using the SuperScript® III Platinum® One-Step Quantitative real-time PCR system (Life Technologies) with the following probe and primers: RSV-A probe CAC CAT CCA ACG GAG CAC AGG AGA T (5’-labeled with 6-FAM and 3’ labeled with BHQ1), RSV-A forward primer AGA TCA ACT TCT GTC ATC CAG CAA and RSV-A reverse TTC TGC ACA TCA TAA TTA GGA G. For initial viral quantification purposes, a commercially available RSV-A2 strain was purchased from (Advanced Biotechnologies Inc). The MOIs used for infectivity experiments were based on genome copies/ml measured by qPCR and the infectious viral titers were measured as TCID_50_ on HEp-2 cells. Briefly, 10-fold dilutions of virus were added onto Hep2 cells for 2 h at 37°C, then inoculum was removed, and fresh maintenance medium was added. The plates were examined in the microscope on day7 and the number of wells for each dilution that was positive for GFP was noted. The titer was calculated according to the method of Karber. The viral stock of 7×10^8^ genome copies/ml corresponded to log_10_ 6.7 TCID_50_/mL.

For HSV-1, strain F HSV-1 stocks were prepared by infecting African green monkey kidney (VERO) cells at 80– 90% confluency in DMEM (Invitrogen) with 5% FBS or in serum-free DMEM. The virus was harvested 2 d after infection. Cells were subjected to two freeze-thaw cycles and spun at 20,000 g for 10 min to remove cell debris. Clarified supernatant was aliquoted and stored at −80°C until use. Non-infected cell medium was prepared with the same procedure without viral infection. Plaque assay was used to determine viral titers. 10-fold dilutions of virus were added onto VERO cells for 1 h at 37°C, then inoculum was removed, and fresh medium containing 0.5% carboxymethyl cellulose (Sigma Aldrich) was added. Cells were fixed and stained 2d later with a 0.1% crystal violet solution and the number of plaques was counted.

### Human BALF samples

The use of human BALF samples for the current study was approved by the Regional Committee for Ethical Review in Stockholm (D. No 2016/1985-32). All donors had given oral and written informed consent to participate in the bronchoscopy study, in line with the Helsinki Declaration. Briefly, healthy subjects of male and female gender were recruited at the Department of Respiratory Medicine and Allergy, Karolinska University Hospital, Solna. These subjects were examined after denying regular tobacco smoking and history of allergy or lung disease during an interview. The final inclusion required that these subjects displayed no signs of pulmonary or disease during clinical examination, spirometry and clinical blood testing including electrolytes, white cell differential counts and C-reactive protein. Bronchoscopy with bronchioalveolar lavage (5 × 50 ml of sterile and phosphate-buffered saline) was performed according to clinical routine at Karolinska University Hospital, Solna. The obtained BALF was concentrated using 5 kDa cutoff 4 ml spin concentrator (Agilent Technologies, USA) before infectivity experiments.

### Lipid vesicle preparation and characterization

1,2-distearoyl-sn-glycero-3-phosphocholine (DSPC), 1,2-dioleoyl-3-trimethylammonium-propane (DOTAP), 1,2-distearoyl-sn-glycero-3-phospho-(1’-rac-glycerol) (DSPG), and cholesterol were bought from Avanti Polar Lipids, Inc. (Alabaster, AL, USA). All other compounds were bought from Sigma-Aldrich (St. Louis, MO, USA). The lipids were dissolved in chloroform in molar ratios of 9:1 (DSPC / cholesterol) for neutral liposomes, 9:1 (DSPC / DOTAP) for cationic liposomes and 9:1 (DSPC / DSPG) for anionic liposomes. The liposomes were formed by the thin film hydration method followed by extrusion through a polycarbonate membrane. Briefly, chloroform was evaporated by heating the sample tube to 65 °C and gradually reducing the pressure to 70 mbar under a nitrogen flow in a vacuum rotary evaporation system (Büchi R-114, Büchi Labortechnik AG, Flawil, Switzerland). The resulting thin lipid layer was hydrated with 500 μL of PBS by gently stirring the tube in a water bath (65 °C) for 1 h. The sample was then extruded 11 times (65 °C) through polycarbonate membrane (pore size of 200 nm) with a syringe extrusion device (Avanti Polar Lipids) after which the sample was quickly cooled down and stored in a refrigerator. The concentration of the samples was 19 µmol/mL. The size of the liposomes was analyzed with a Zetasizer APS dynamic light scattering automated plate sampler (Malvern Instruments, Malvern, United Kingdom). The concentration of liposomes was determined using Nanosight (Malvern, USA). The zeta potential was measured at 25 °C in DTS 1070 folded capillary cells with Zetasizer Nano ZS (Malvern Instruments).

### Protein corona proteomics

FBS was obtained commercially (ThermoFisher, USA). HP was obtained and pooled from at least 3 different healthy donors. jHP was obtained and pooled from at least 3 different infants participating in a prospective cohort of 281 children born into the cohort between 1997 and 2000 in Stockholm, Sweden. The study was approved by the Human Ethics Committee at Huddinge University Hospital, Stockholm (Dnr 75/97). MP was obtained and pooled from at least three different Indian rhesus macaques that were RSV seronegative. Ethical permit Dnr N2 / 15, Department of Medicine, Karolinska University Hospital, Solna. All protein corona experiments described below were performed in technical triplicates. For the viral corona proteomic experiments, freshly-harvested serum-free RSV supernatant (not-sucrose cushion concentrated) containing 6.6 × 10^9^ RSV genome equivalents or serum-free DMEM produced HSV-1 stocks (2.1 × 10^8^ PFU/ml) was incubated with 10% v/v solutions of different biological fluids in for 1h at 37 °C. Before adding the viruses, all the biological fluid solutions were adjusted to 10 mM sodium citrate (SigmaAldrich, Germany) to prevent coagulation. As a control, uninfected cell supernatant prepared in a similar way was also incubated with the biological fluid solutions. For lipid vesicles, 3 × 10^11^ of 200 nm positively or negatively charged lipid vesicles were incubated with 10% solutions of biological fluids at similar conditions. After incubation, viral/nanoparticle corona complexes were spun at 20,000 g at 4 °C for 1 h, supernatant removed, and the pellet resuspended in 1 ml PBS. The pellet was washed twice with PBS using the same centrifugation conditions then boiled at 95 °C for 5 min. before measuring the protein content using Micro BCA™ protein assay kit (ThermoFisher, USA). The viral/nanoparticle corona complexes were then resuspended in PBS and adjusted with ammonium bicarbonate to a final concentration of 20 mM. The samples were reduced by addition of 1:20 v/v 100 mM dithiothreitol (DTT) for 45 min. at 56 °C and alkylated with 1:20 v/v 100 mM iodoacetamide (IAA) for 45 min at RT in the dark. Proteins were digested with trypsin (MS gold, Promega) 0,1 µg/µl overnight at 37 °C (trypsin : protein ratio 1:50). For sample clean-up, the samples were applied to strong cation exchange SCX microcolumns (Strata-XC Phenomenex). The microcolumns were initially washed with 100% methanol followed by MilliQ grade water. The samples were adjusted to >0.1% formic acid and then applied to the columns. After washing with 30% methanol and 0.1% formic acid the samples were eluted with 30% methanol and 5% ammonium hydroxide. Samples were dried in a SpeedVac and submitted to mass spectrometry (MS). For MS analysis, samples were dissolved in 3% ACN, 0.1% formic acid to end-concentration 1µg/µL. 1-2 µl of sample were analyzed with a hybrid LTQ-Orbitrap Velos or with an Orbitrap Fusion Tribrid Mass Spectrometer (ThermoFisher, USA). For the Velos system, an Agilent HPLC 1200 system (Agilent) was used to provide the 70 min gradient. Data-dependent MS/MS (centroid mode) followed in two stages: first, the top-5 ions from the master scan were selected for collision induced dissociation with detection in the ion trap (ITMS); and then, the same 5 ions underwent higher energy collision dissociation (HCD) with detection in the Orbitrap (FTMS). For the Fusion system, An UltiMate 3000 UHPLC System was used to provide the 70 min gradient. Data-dependent MS/MS (centroid mode) followed in two stages: first, the top-10 ions from the master scan were selected for collision-induced dissociation with detection in the ion trap (ITMS); and then, the same 10 ions underwent higher energy collision dissociation (HCD) with detection in the Orbitrap (FTMS). The data was searched by Sequest under the Proteome Discoverer 1.4.1.1.4 software (ThermoFisher) against the following Uniprot protein sequence databases; bos taurus version 170310, rhesus macaque version 170704, human version 160210, RSV version 160210 HHV1 version 180613, and Chlorocebus sabaeus version 180613 using a 1% peptide false discovery rate (FDR) cut-off limit. For samples containing proteins from multiple species, combined species databases were created by merging the corresponding protein databases. For label-free quantification, protein MS1 precursor area was calculated as the average of the top-three most intense peptides. Only proteins significantly detected (FDR 1%) in all three technical replicates in each sample were used in the downstream data analysis.

### Data analysis

Only proteins significantly detected (FDR 1%) in all three technical replicates in each sample were used in the data analysis. PCA analysis and hierarchical clustering of the filtered proteomics data (described above) was performed using ClustVis (https://biit.cs.ut.ee/clustvis_large/). Gene ontology gene list enrichment analysis was performed using ToppFun (https://toppgene.cchmc.org/enrichment.jsp). Average coefficient of variation (CV) was calculated for each sample based on protein precursor area. Spearman correlation matrices were performed using Morpheus software based on precursor area of proteins significantly detected (FDR 1%) in all three technical replicates in each sample (https://software.broadinstitute.org/morpheus). For other experiments, non-parametric Kruskal-Wallis unpaired test was used followed by Mann-Whitney test or Dunn’s post-test to compare the data. Six replicates from two independent experiments. Statistics were calculated using GraphPad Prism 7 software.

### ELISA

The detection of specific anti-RSV IgG antibodies in biological fluids was performed using Human Anti-Respiratory syncytial virus IgG ELISA Kit (ab108765, abcam®, Sweden), according to manufacturer’s protocol. All biological fluids (FBS, HP, MP, BALF) were diluted to a protein concentration of 0.3 mg/ml before performing the assay and results were compared to the positive, cutoff, and negative controls provided by the kit manufacturer.

### RSV infectivity

HEp-2 cells were seeded in maintenance medium until they reached 50-60% confluency. On the day of infection, the cells were washed twice with warm PBS and medium was changed to VP-SFM with gentamycin. Before adding to the cells, sucrose cushion-concentrated RSV stocks were pre-incubated with different biological fluids (FBS, HP, MP, BALF) at a final protein concentration of 0.3 mg/ml in VP-SFM with 10 mM sodium citrate for 1 h at 37 °C. The corona-precoated viruses were then added to the cells in serum-free conditions (diluted 10x) at a MOI of 1. 24h post-infection, the medium was changed back to growth medium and cells were visualized with fluorescence microscopy 72h post-infection. After visualization, the cells were washed and stained with LIVE/DEAD™ Fixable Far Red Dead Cell Stain Kit (ThermoFisher, USA) for 30 min at 4 °C in Dulbecco’s phosphate-buffered saline (DPBS no calcium, no magnesium, ThermoFisher), then fixed and washed for flow cytometry using Cytofix/Cytoperm™ (BD, USA) according to manufacturer’s protocol. Cells were then scraped and resuspended in DPBS, and the data was acquired using MACSQuant® Analyser 10 flowcytometer (Miltenyi Biotec, Sweden). The data was analyzed by FlowJo software (TreeStar) by excluding the dead cells stained with far-red fluorescent dye and subsequent calculation of GFP positive cells within the viable cell population. For experiments with palivizumab, cells were treated with different concentrations of the antibody before infection.

### MoDC differentiation

Human monocytes were negatively selected from buffy coats using the RosetteSep Monocyte Enrichment Kit (1mL/10mL buffy coat; StemCell Technologies) and differentiated into moDC, using GM-CSF (250ng/mL; PeproTech) and IL-4 (6.5ng/mL; R&D Systems) for 6 days in 37°C, 5% CO_2_ at a density of 5×10^5^ cells/mL in RPMI 1640 completed with 10% FCS, 1mM sodium pyruvate, 10mM HEPES, 2mM L-glutamine, and 1% and penicillin/streptomycin (ThermoFisher, USA). Immature moDC were exposed to RSV pre-incubated with different biological fluids for 4h in serum free media, washed and then incubated in serum containing medium for 72 h before analyses of CD86 and RSV-GFP expression. Dead cells were excluded using Live/Dead fixable near-IR dead cell stain kit (ThermoFisher). Flow cytometry sample data were acquired on a Fortessa (BD Biosciences) and the analysis was performed in FlowJo software (TreeStar).

### Thioflavin-T Assay

NNFGAIL, GNNQQNY, Aβ_42_, Aβ_40_ were synthesized and purified by Pepscan (The Netherlands) and the final molecular weight was verified by MS. 10 µl of dimethyl sulfoxide (DMSO) (ThermoFisher, USA) were added to 1 mg aliquots of the peptide and running stocks were prepared in MQ water. ThT (Sigma Aldrich) was prepared at 4 mM in MQ water. For the assay with RSV and NNFGAIL or GNNQQNY, 50µl of 1mM peptide were incubated with 150µl of 4 mM ThT solution and 100 µl of RSV (freshly-harvested serum-free RSV supernatant, not-sucrose cushion concentrated, 3 × 10^8^ genome copy/ml), non-infected supernatant, serum-free viral production medium (VP-SF medium), VP-SF medium + 5% FBS or lipid vesicles (positively and negatively charged, dimeter= 200 nm, concentration = 1 × 10^10^ particles/ml in VP-SF medium). For the assay with HSV-1 and Aβ_42_ or Aβ_40_, 50 µl of 50 µM peptide were incubated 150 µl of 4 mM ThT solution and 100 µl of HSV-1 (2 × 10^8^ PFU/ml), non-infected supernatant, DMEM serum free medium, DMEM serum free medium + 5% FBS or lipid vesicles (positively and negatively charged, dimeter= 200 nm, concentration = 1 × 10^10^ particles/ml in DMEM serum free medium). ThT fluorescence was measured at 440 nm excitation and 480 nm emission in a black clear-bottom 96- well plates (Corning, USA) at 440 nm excitation and 480 nm emission at 10-15 min. intervals (from bottom with periodic shaking) over 12-24 h on SpectraMax i3 microplate reader (Molecular Devices, USA). Curves were fitted using GraphPad Prism software.

### Electron microscopy

For cell sections with RSV, HEp-2 cells were seeded in 6 cm dishes in maintenance medium until 70-80% confluent, then washed and medium replaced with VP-SFM with gentamycin before infecting with RSV at MOI 100 in serum-free conditions or in 50% v/v of different biological fluids. Cells were then fixed with 2.5 % glutaraldehyde in 0.1M phosphate buffer, pH 7.4 at room temperature for 30 min. The cells were scraped off and transferred to an Eppendorf tube and further fixed overnight in the refrigerator. After fixation, cells were rinsed in 0.1M phosphate buffer and centrifuged (100g for 5 min.). The pellets were then fixed with 2% osmium tetroxide (TAAB, Berks, England) in 0.1M phosphate buffer, pH 7.4 at 4 °C for 2 h, then dehydrated in ethanol followed by acetone and embedded in LX-112 (Ladd, Burlington, Vermont, USA). Ultrathin sections (approximately 50-60 nm) were cut by a Leica EM UC 6 (Leica, Wien, Austria). Sections were stained with uranyl acetate followed by lead citrate and imaged in a Tecnai 12 Spirit Bio TWIN transmission electron microscope (FEI Company, Eindhoven, The Netherlands) at 100 kV. Digital images were captured by a Veleta camera (Olympus Soft Imaging Solutions, GmbH, Münster, Germany). For cryoimmunoelectron microscopy (iEM), cells were fixed in 3 % paraformaldehyde in 0.1 M phosphate buffer. Samples were then rinsed with 0.1M phosphate buffer and infiltrated in 10% gelatin. Then the specimens were infiltrated into 2.3 M sucrose and frozen in liquid nitrogen. Sectioning was performed at −95°C and mounted on carbon-reinforced formvar-coated, 50 mesh Nickel grids. Immunolabelling was performed as follows: grids were placed directly on drops of 2% normal goat serum (DAKO, Glostrup, Denmark) in 0.1 M phosphate buffer to block non-specific binding then incubated with the primary antibodies: mouse anti-RSV fusion protein monoclonal antibody (MAB8599, Millipore, USA) or goat anti-human IgG (Licor, USA) or mouse anti-surfactant protein A antibody (abcam, ab51891). Antibodies were diluted 1:50 in 0.1M of phosphate buffer containing 0.1% normal goat serum overnight in a humidified chamber at room temperature. The sections were washed using the same buffer and bound antibodies were detected using secondary antibodies coated with 10 nm gold (BBI solution, Analytic standard, Sweden) at a final dilution of 1:100. Sections were then rinsed in buffer and fixed in 2% glutaraldehyde, stained with 0,05% uranyl acetate, embedded in 1% methylcellulose and then examined in a Tecnai G2 Bio TWIN (FEI company, Eindhoven, The Netherlands) at 100 kV. Digital images were captured by a Veleta camera (Soft Imaging System GmbH, Műnster, Germany). For TEM of viruses with amyloids, 100 µl of RSV (3 × 10^8^ genome copies/ml) or HSV-1 (2 × 10^7^ PFU/ml) were incubated with 50µl 1mM NNFGAIL (for RSV) or 50µM Aβ_42_ (for HSV-1) for 100 min. at 37 °C. Samples were applied to Formvar/carbon coated 200 mesh nickel grids (Agar Scientific, UK), then negatively stained using an aqueous solution of uranyl acetate (1%) and visualized.

### Western blotting

Serum-free produced RSV was incubated with BALF or HP both at a protein concentration of 0.1 mg/ml at a ratio of 1:1 v/v (in citrate adjusted conditions to prevent coagulation) for 1h at 37 ℃, then harvested and washed similar to the procedure for corona proteomic analysis. Samples were lysed using a RIPA lysis buffer with protease inhibitors and protein content was measured. Samples were heated with loading buffer to 95°C for 5 min and 1 µg of protein was loaded onto NuPAGE® Bis-Tris 4–12% gels then separated at 80 V for 15 min followed by 100V for 135 min at room temperature using 1x NuPAGE® MES SDS running buffer (Invitrogen). The gels were then transferred onto nitrocellulose membranes using the IBlot® system (Invitrogen) and subsequently the membranes were blocked with Odyssey Blocking Buffer (LI-COR Biosciences GmbH) for 1.5 h. Membranes were incubated overnight at 4 ℃ with anti-Surfactant Protein A antibody [6F10] (ab51891, abcam) diluted at 1:500 then probed the next day using goat anti-human IgG (IRDye 800, red, LI-COR Biosciences) and goat anti-mouse IgG (IRDye 680, green, LI-COR Biosciences). To normalize the bands to a loading control, scanned membranes were stripped for 10 min at 65 °C using 1x New Blot Nitro Stripping buffer (LI-COR Biosciences), blocked for 1.5 h, and then incubated with anti-respiratory syncytial virus antibody (ab20745, abcam)^65^ at 1:500 dilution for 2h at RT before probing with secondary antibody (donkey anti-goat, IRDye 680, green, LI-COR Biosciences). All western blot signals were scanned using the Odyssey Imager (LI-COR Biosciences GmbH) and quantification was performed on the images using ImageJ software.

### Animal experiments

To evaluate the impact of the HSV-1 on brain β-amyloid (Aβ) levels, 3-month-old transgenic 5XFAD mice (purchased from Jackson Laboratories, Bar Harbor, Maine, US) were randomly divided into 2 groups. The animals were injected either with the HSV virus or non-infected supernatant using a micro-infusion pump (The Harvard Apparatus Pump Series, Harvard Bioscience, USA) into the right lateral ventricles. Briefly, surgical anesthesia was induced with 5% isoflurane and maintained with 1.8% isoflurane (in 30% O_2_/70% N_2_O). The temperature of the animals was maintained at 37 +/- 0.5 °C using a thermostatically controlled heating blanket connected to a rectal probe (PanLab, Harvard Apparatus, Barcelona, Spain). The skin was opened and the scull exposed. A small hole approximately 1 mm in diameter was drilled into the following coordinates: m/l (medial/lateral) +1.1 mm, a/p (anterior/posterior) −0.3 mm, d/v (dorsal/ventral) −2.0 mm. The mice were infused with either 10 µl of HSV- 1 virus (2 × 10^7^ PFU/ml) or with non-infected supernatant as control. After injection the wound was sutured and the animals placed in individual cages to recover for 48h. All animal work was approved by the Animal Care and Use Committee of the University of Eastern Finland (Kuopio) and performed according to the guidelines of National Institutes of Health for animal care. Mice were euthanized at 48 h after ICV (intracerebroventricular) infusion for tissue collection. The mice were anesthetized with an overdose of Avertin followed by transcardial perfusion with heparinized saline (2500 IU/L). The ICV infused brain hemisphere were removed and post-fixed in 4% PFA followed by cryoprotection in 30% sucrose. The hemibrains were frozen in liquid nitrogen and cryosectioned into 20-μm sagittal sections and stored in the anti-freeze solution. Six consecutive sagittal brain sections at 400 μm intervals were selected for immunohistological staining from each mouse. Aβ deposits were detected using primary anti-body specific to isoforms Aβ 37-42 (Aβ D54D2 XP, Cell Signaling Technology, 1:100 dilution, overnight at RT) and further visualized by fluorescent Alexa 568 secondary antibody (1:500 dilution, ThermoFisher Scientific). For quantification of Aβ immunoreactivities, the stained sections were imaged using 10x magnification in Zeiss Axio ImagerM.2 microscope equipped with Axiocam 506 mono CCD camera (Carl Zeiss, Oberkochen, Germany) running ZEN software (Carl Zeiss) for tailing and stitching of the images. Cortical and hippocampal Aβ immunoreactivities were quantified from 6 stained sections at 400 um intervals per animal using MatLab (MathWorks, MatLab 2017b). The accuracy of the analysis was confirmed by re-analyzing the images using ImageJ 1.50i software.

### Data Availability

All data generated or analyzed during this study are included in this published article and its supplementary information files.

## Acknowledgements

We would like to acknowledge the Svenska Sällskapet för Medicinsk Forskning (SSMF) for supporting Kariem Ezzat, Swedish Research Council (K2015-99X-22880-01-6) and Stockholm University for supporting Anna-Lena Spetz, Vetenskapsrådet, SSMF, and the Swedish foundation for Strategic Research for supporting Samir EL-Andaloussi, Dr. Michael N. Teng at the University of South Florida for providing the RSV-GFP virus, Professor Arto Urtti at the University of Helsinki for the lipid vesicles, the Instumentarium Science Foundation for supporting Otto K. Kari, the service of the Electron Tomography Facility at Karolinska Institutet, and the Academy of Finland for supporting the research of Tarja Malm. Anders Lindén obtained project funding from the Swedish Heart-Lung Foundation (No. 20150303), the Swedish Research Council (No. 2016-01653), as well as federal funding from Karolinska Institutet and through the Regional Agreement on Medical Training and Clinical Research (ALF, No. 20140309) between Stockholm County Council and Karolinska Institutet. No funding was obtained from the tobacco industry.

## Author Contributions

K.E. conceived the concept, designed, performed and analyzed experiments and wrote the paper. M.P. designed, conducted and analyzed proteomics experiments. S.P. conducted and analyzed moDC experiments and FACS data. T.R. analyzed proteomics data. P.J. designed, performed and analyzed moDC experiments. A.D. conducted initial RSV propagation. B.B. performed and analyzed FACS experiments. M.J.S. and J.K.S provided HSV-1 stocks. B.L, M.S. and A.L. provided BALF samples. E.A.T provided MP samples. O.S. performed and analyzed western blotting experiments. O.K.K and T.L. synthesized, characterized and provided lipid vesicles. E.S.E and C.N. provided jHP samples. Y.I. and T.M. designed, performed and analyzed animal experiments. S.M. performed electron microscopy experiments. M.J.A.W, U.F.P., J.L. and S.E.A. provided critical feedback. A-L.S. designed and analyzed experiments and provided critical feedback. All authors read and commented on the manuscript.

