## Supplementary Figures for "The Viral Protein Corona Directs Viral Pathogenesis and Amyloid Aggregation"

**Ezzat K. et al.**

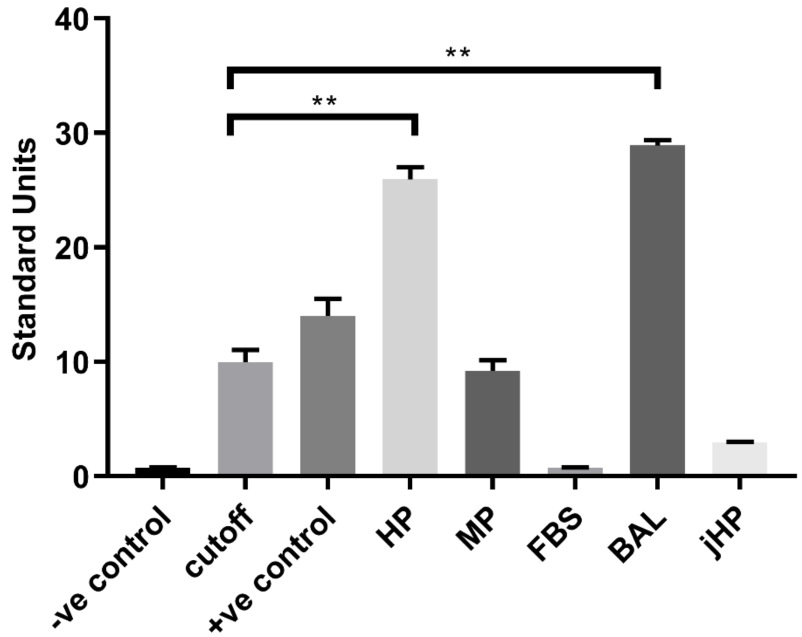


**Supplementary Figure 1. ELISA analysis of anti-RSV IgG antibodies in different biological fluids.** Different biological fluids were diluted to a protein concentration of 0.3 mg/ml, and then ELISA was used to quantify RSV specific IgG antibodies. Significant differences were assessed by Mann-Whitney test and are indicated by ***P* <.01. Data are shown as means ± SEM of six replicates from two separate experiments.

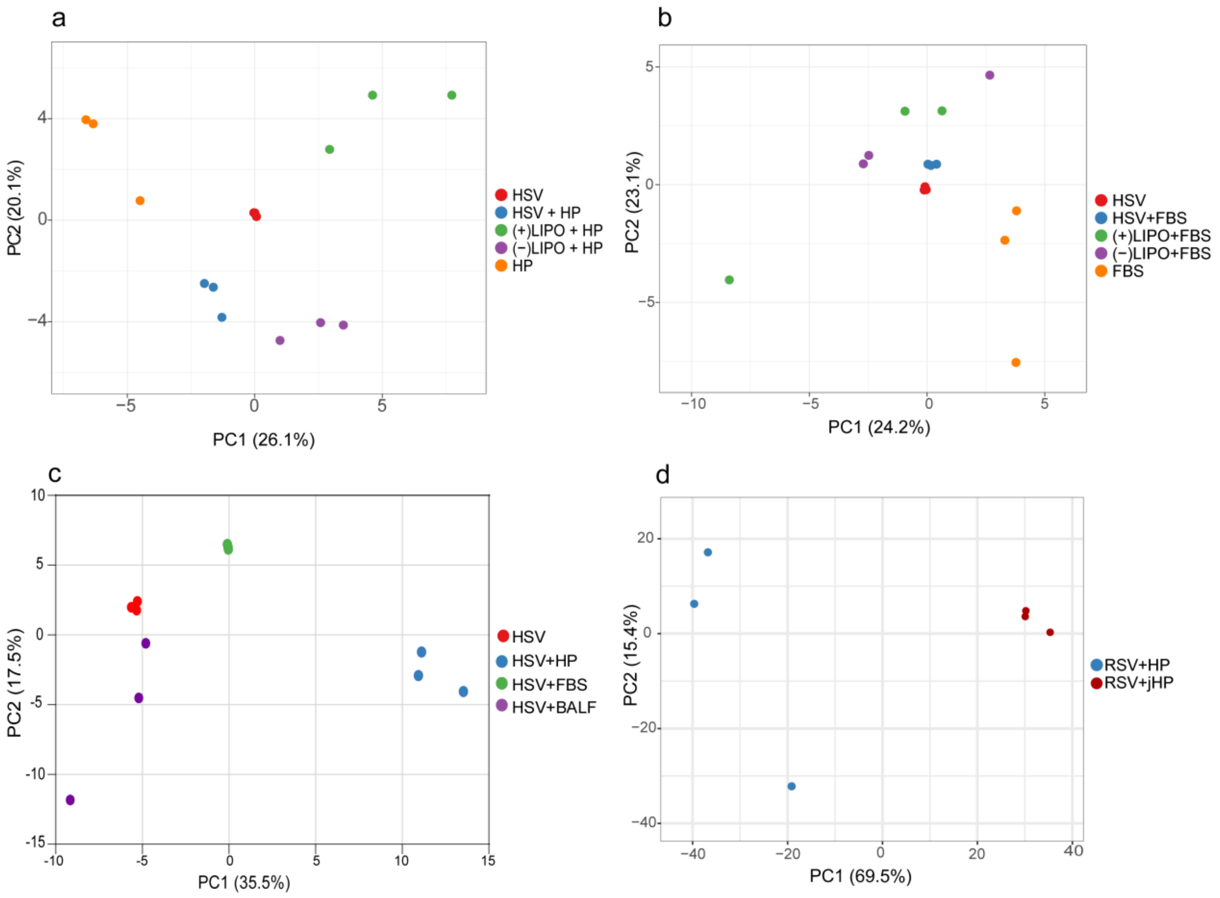


**Supplementary Figure 2. Principal component analyses (PCA) of the corona proteomic profiles**. Triplicate samples were incubated with 10% solutions of different biological fluids for 1h at 37 ˚C, then re-harvested, washed and finally analyzed by MS. (-)Lipo = negatively charged lipid vesicles, 200 nm, (+)Lipo = positively charged lipid vesicles, 200 nm. Only proteins significantly detected (FDR 1%) in all three replicates in each condition were used. (**a**) PCA comparing proteomic profiles of HSV-1 and controls in human plasma (HP) (**b**) PCA comparing proteomic profiles of HSV-1 and controls in fetal bovine serum (FBS). (**c**) PCA comparing proteomic profiles of HSV-1 in different biological fluids. (**d**) PCA comparing corona proteomic profiles of RSV in adult human plasma (HP) vs. juvenile human plasma (jHP).

a

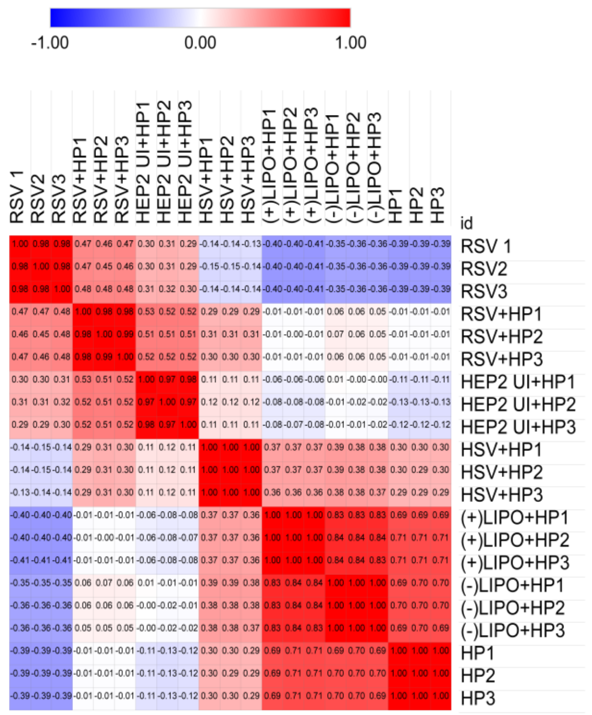


b

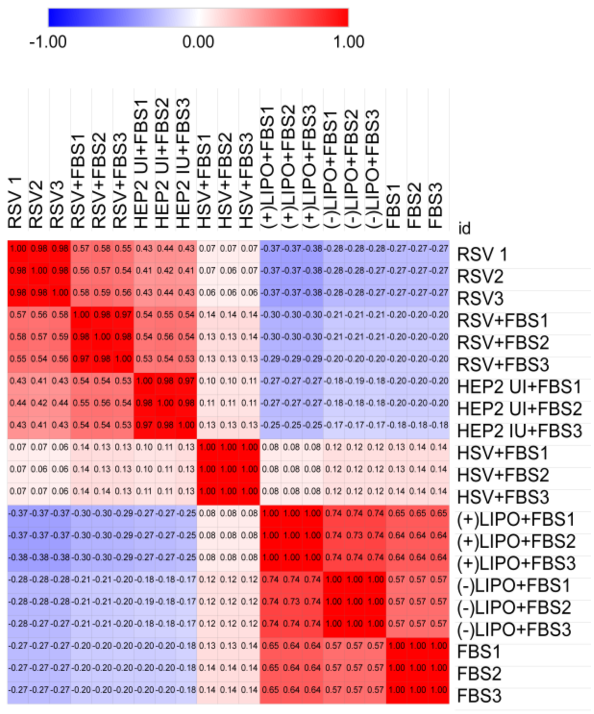


c

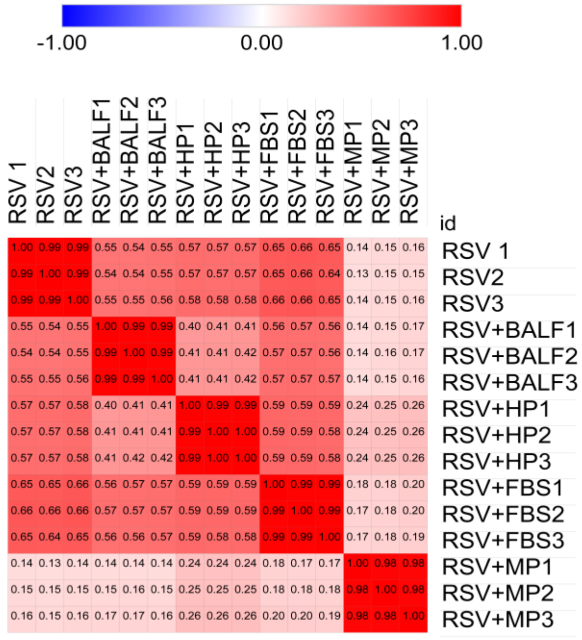


d

| MATRIX | CV% |
|---|---|
| FBS | 29 |
| HP | 19 |
| (-)Lipo + FBS | 42 |
| (-)Lipo + HP | 36 |
| (+)Lipo + FBS | 41 |
| (+)Lipo + HP | 28 |
| HSV+FBS | 16 |
| HSV+HP | 23 |
| RSV+BALF | 33 |
| RSV+MP | 31 |
| NI+FBS | 31 |
| NI+HP | 22 |
| RSV+FBS | 25 |
| RSV+HP | 26 |
| RSV | 33 |
| HSV | 19 |
| RSV+jHP | 14 |

**Supplementary Figure 3. Correlation matrices for each of the PCAs presented in Figure 1.** (**a-c**) Spearman correlation coefficients between each of the samples in the PCA based on precursor area of proteins significantly detected (FDR 1%) in all three replicates, (**a**) for samples included in Fig. 1A, (**b**) For samples included in Fig. 1B and (**c**) for samples included in Fig 1C. (**d**) Average coefficients of variation (CV) calculated for replicates in each corona condition based on protein precursor area.

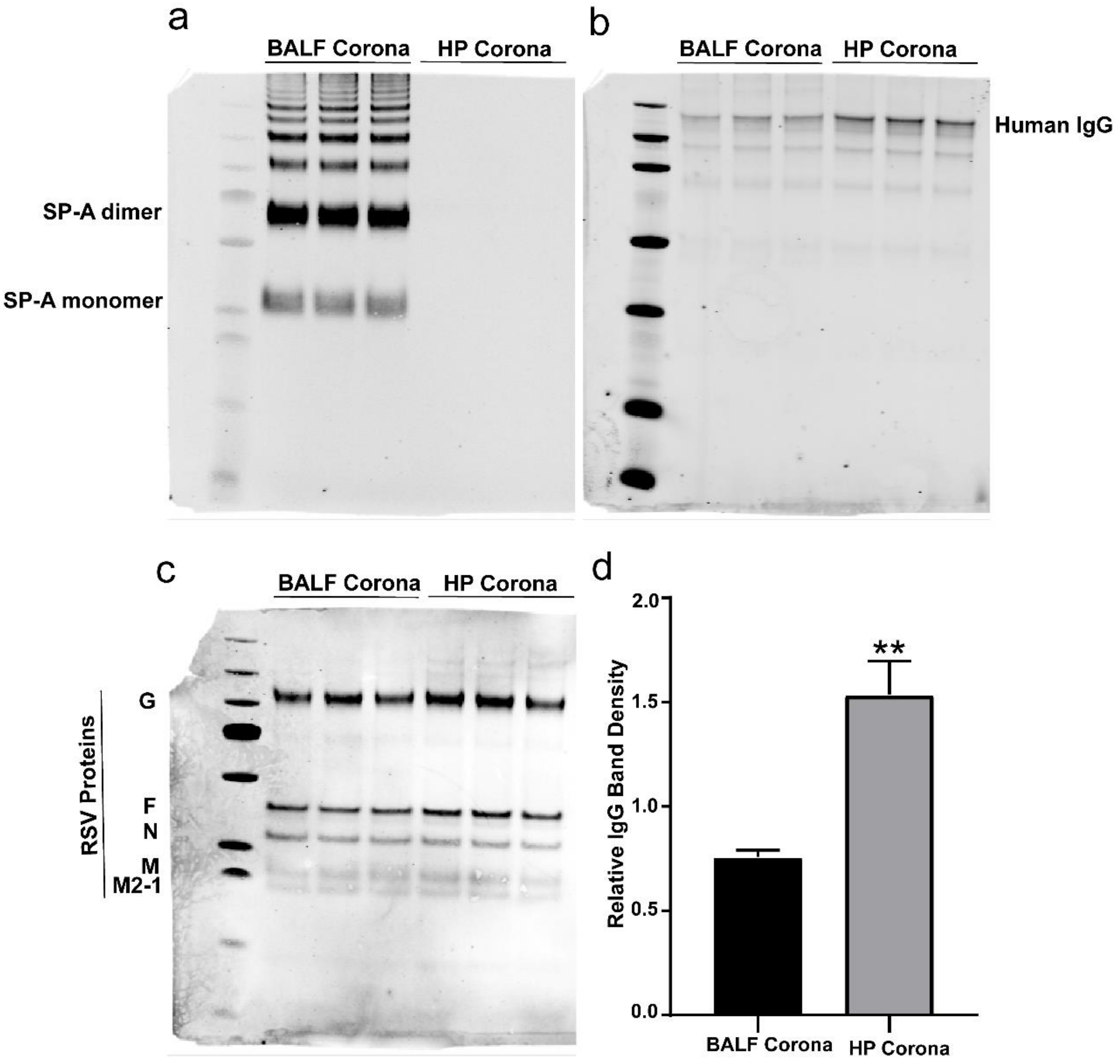


**Supplementary Figure 4. Detection of specific corona proteins using western blot.** Serum-free produced RSV was incubated with BALF or HP at a protein concentration of 0.1 mg/ml at a ratio of 1:1 v/v for 1h at 37 °C then harvested and washed in a procedure similar to the corona proteomic analysis. (**a**) Bands detected using anti SP-A antibody appear only in the BALF corona samples showing SP-A bands together with higher molecular weight SP-A containing complexes. No SP-A bands were detected in the HP corona samples. (**b**) Bands detected using anti-human IgG antibody appear in both BALF and HP corona, with higher intensity in the latter. (**c**) RSV viral proteins were used as loading controls. (**d**) IgG band intensity was quantified and normalized to the G protein loading control band intensity using ImageJ software. Data is represented as mean relative intensity ±SEM and significance was calculated using Student's T-test and is indicated by **P <.01.

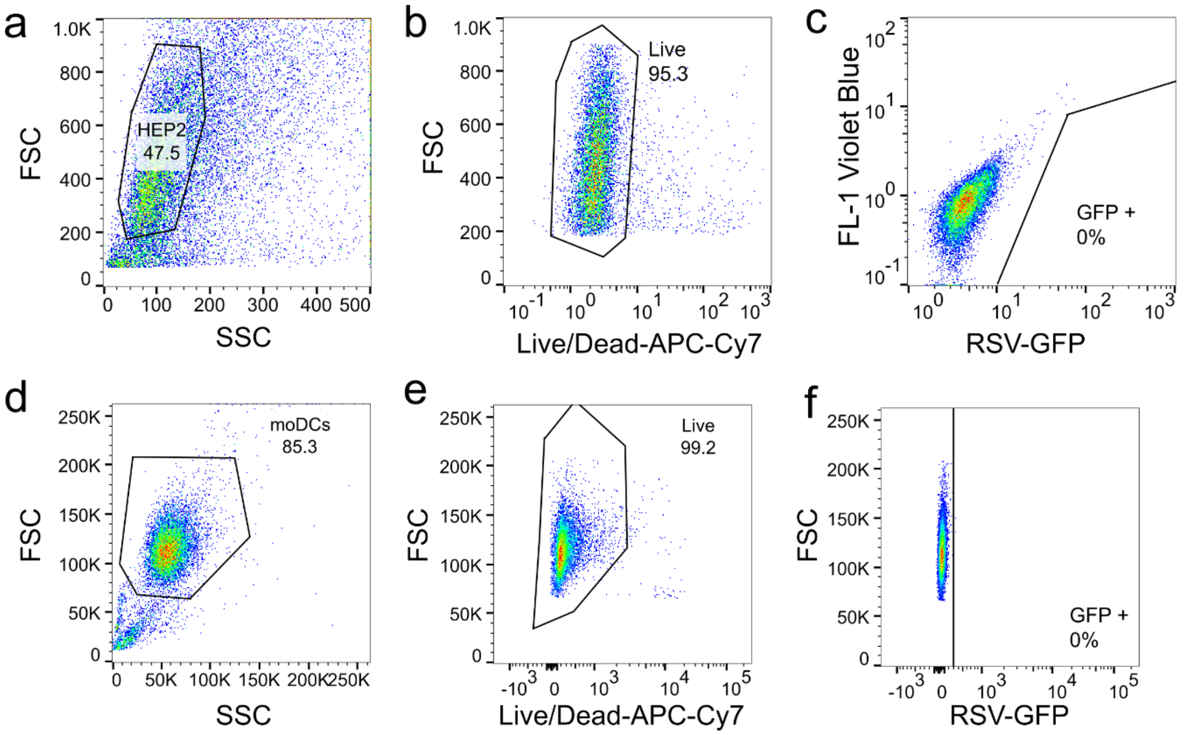


**Supplementary Figure 5. Methodology of FACS gating** (**a-c**) Uninfected HEp-2 cells and (**d-f**) MoDCs. Cells were first gated based on FSC/SSC (**a** and **d**), followed by gating on living cells (**b** and **e**) and lastly the GFP gate (**c** and **f**). Autofluorescent, non-infected Hep-2 cells were excluded using the FL1 channel.

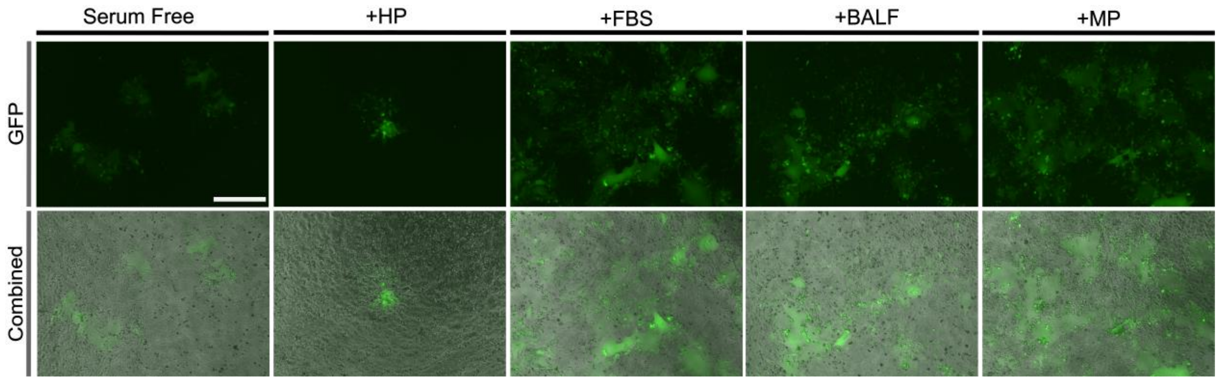


**Supplementary Figure 6. Fluorescence microscopy of RSV infectivity.** Representative pictures showing infectivity and syncytia formation in in HEp-2 cells using virus stocks in serum-free conditions or with different coronae resulting from pre-incubation with HP, FBS, BALF or MP (n=3). The upper row presents images the fluorescence channel and the lower row represents images of combined fluorescence and phase contrast channels. Bar= 500µM.

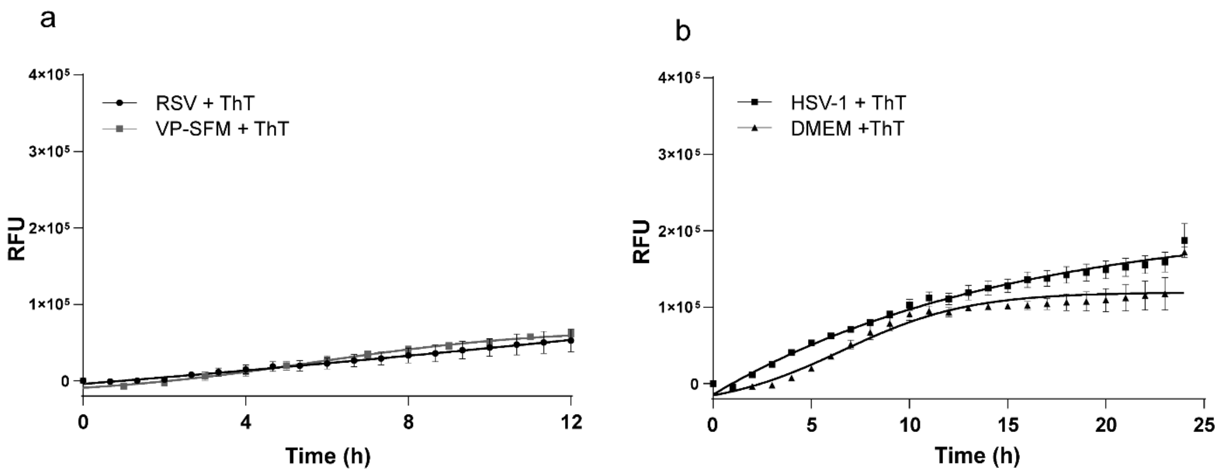


**Supplementary Figure 7. Curves representing the background interaction of ThT with the growth media in absence of peptides** (**a**) for RSV and (**b**) for HSV-1. Means ±SEM of six replicates from two separate experiments are shown.
